# Improved Characterization of the Solution Kinetics and Thermodynamics of Biotin, Biocytin and HABA Binding to Avidin and Streptavidin

**DOI:** 10.1101/410548

**Authors:** Roberto F. Delgadillo, Timothy C. Mueser, Kathia Zaleta-Rivera, Katie A. Carnes, José González-Valdez, Lawrence J. Parkhurst

**Affiliations:** Department of Chemistry, University of Nebraska - Lincoln, Lincoln, NE 68588-0304, USA; Tecnologico de Monterrey, School of Engineering and Sciences, Av. Eugenio Garza Sada 2501 Sur, Monterrey, NL 64849, Monterrey, Mexico; Department of Chemistry and Biochemistry, University of Toledo, Toledo, OH, 43606, USA; Department of Bioengineering, University of California San Diego, San Diego, CA, 92093-0412, USA.; GlaxoSmithKline, Biopharmaceutical Analytical Sciences Department. King of Prussia, PA, 19406, USA.

## Abstract

The high affinity (K_D_ ∼ 10^−15^ M) of biotin to avidin and streptavidin is the essential component in a multitude of bioassays with many experiments using biotin modifications to invoke coupling. Equilibration times suggested for these assays assume that the association rate constant (k_on_) is approximately diffusion limited (10^9^ M^−1^s^−1^) but recent single molecule and surface binding studies indicate they are slower than expected (10^5^ to 10^7^ M^−1^s^−1^). In this study, we asked whether these reactions in solution are diffusion controlled, what reaction model and thermodynamic cycle described the complex formation, and the functional differences between avidin and streptavidin. We have studied the biotin association by two stopped-flow methodologies using labeled and unlabeled probes: I) fluorescent probes attached to biotin and biocytin; and II) unlabeled biotin and HABA, 2-(4’-hydroxyazobenzene)-benzoic acid. Native avidin and streptavidin are homo-tetrameric and the association data show no cooperativity between the binding sites. The k_on_ values of streptavidin are faster than avidin but slower than expected for a diffusion limited reaction in both complexes. Moreover, the Arrhenius plots of the k_on_ values revealed strong temperature dependence with large activation energies (6-15 kcal/mol) that do not correspond to a diffusion limited process (3-4 kcal/mol). The data suggest that the avidin binding sites are deeper and less accessible than those of streptavidin. Accordingly, we propose a simple reaction model with a single transition state for non-immobilized reactants whose forward thermodynamic parameters complete the thermodynamic cycle in agreement with previously reported studies. Our new understanding and description of the kinetics, thermodynamics and spectroscopic parameters for these complexes will help to improve purification efficiencies, molecule detection, and drug screening assays or find new applications.

## 1. INTRODUCTION

The extremely high affinity of biotin (B_7_, vitamin H) for avidin (AV) and streptavidin (SAV) is widely exploited in biotechnology and biochemistry in a vast array of applications.^1,2^ It has been used in molecular biology as markers to identify functional moieties in proteins, receptors^3^ and the development of bioprocessing affinity chromatography columns for the recovery of highly valued biomolecules.^4^ More recently, advances in the characterization of these complexes have allowed the development of highly specific immunoassays, biosensors, and “omic” tools for disease identification and its molecular mechanism elucidation.^5–8^ Furthermore, B_7_ and avidin-like interactions can be exploited for imaging purposes in the development of assays in vivo and real-time visualization of intracellular or other type of biological processes^9,10^ and nanoscale drug delivery systems of small molecules, proteins, vaccines, monoclonal antibodies, and nucleic acids.^11^ SAV and B_7_ are used in a Fluorescence Resonance Energy Transfer (FRET)^12^ systems for drug High Throughput Screening (HTS) applications, commercially know as Homogeneous Time-Resolved Fluorescence (HTRF).^13–15^ Furthermore, it has been suggested that these proteins function in nature as antimicrobial agents by depleting B_7_ or sequestering bacterial and viral DNA,^16,17^ and questions concerning their biological importance increase as new avidin-like proteins are discovered. For example, rhizavidin was discovered from proteobacterium *Rhizobium etli*,^18,19^ tamavidin from the basidiomycete fungus *Pleurotus cornucopiae*,^20^ xenavidin from the frog *Xenopus tropicalis*,^21^ bradavidin from *Bradyrhizobium japonicum*,^22,23^ and other AV related proteins have been isolated from chicken, *Gallus gallus*.^24–28^

The monomers of AV and SAV are eight stranded anti-parallel beta-barrels with several aromatic residues forming the biotin binding site at one end of the barrel.^29^ Two monomers lie parallel to each other forming a dimer with an extensive interface and two dimers associate forming the weaker interface of the homotetramer. The apo-tetramer has modest thermal stability and the protein becomes highly thermal stable with ligand bound.^30^ Intriguingly, the dimeric interface appears to be necessary for high affinity. Introducing two interface mutations, Trp110 to Lys to form dimers of high affinity plus Asn54 to Ala to form monomers, that remains monomeric with ligand bound, that have a significantly reduced affinity (K_D_ ∼ 10^−7^ M). ^31^ The use of monomeric avidin in affinity chromatography allows for reversible binding.

As it can be inferred, new applications for B_7_:avidin related complexes will surely continue to emerge as more derivatives are characterized. However, to obtain reliable and sensitive applications; a better understanding of the thermodynamics, fluorescence behavior of the attached probes, kinetic reaction mechanisms of association and dissociation of B_7_ and avidin-like systems are surely need, which can used to improve purification efficacies, detection, drug screening assays, and develop new nanotechnological applications. Therefore, we want to provide a more global description of the AV-B and SAV-B systems for bio- and nano-technological applications.

The association rate constant of B_7_ binding to AV (k_on_) is assumed to be near diffusion limited since it was first measured by Green^32^ (7.0 × 10^7^ M^−1^s^−1^, pH 5 and 25 °C) employing a quenching experiment that required the quantification, by chromatographic separation, of un-reacted ^14^C-biotin. Since then several widely varying k_on_ values have been reported for both AV and SAV ranging from 1 × 10^5^ M^−1^s^−1^ to 2 × 10^8^ M^−1^s^−1 20,33–36^ with error ranges below 10%.

Despite this information, the kinetic and thermodynamic parameters of the B_7_ association to these AV and SAV proteins have not been studied in systematic detail. Consequently, for this study, *we asked whether the association rate constants (k*_*on*_*) for* B_7_ *binding to AV and SAV are actually diffusion controlled, what are the association model and thermodynamic cycle that describe the reaction process, and the functional differences between AV and SAV.* We analyzed the k_on_ for B_7_ binding to AV and SAV by two stopped-flow (SF) methodologies employing fluorescent dye labeled-and unlabeled-B derivatives. In the first case, the association reactions were monitored with two sensing modalities: Fluorescence change, F(t), and corrected fluorescence anisotropy, rF(t), under pseudo-first-order conditions as a function of temperature, concentration, and pH with the help of three dye-labeled B_7_ probes: 1) Biotin-4-fluorescein (BFl), 2) Oregon green 488 ® biocytin (BcO), and 3) biotin-DNA_ds_*Fl-3’ (Figure 1). The functional cofactor form of B_7_ is biocytin (Bc) which is formed through an amide linkage between the ε-amine of lysine and carboxyl group of biotin. Modified BcO contains a significantly longer linker with respect to BFl which allows analysis of a potential steric effect in the association process, as has been reported elsewhere.^37^

**Figure 1.**
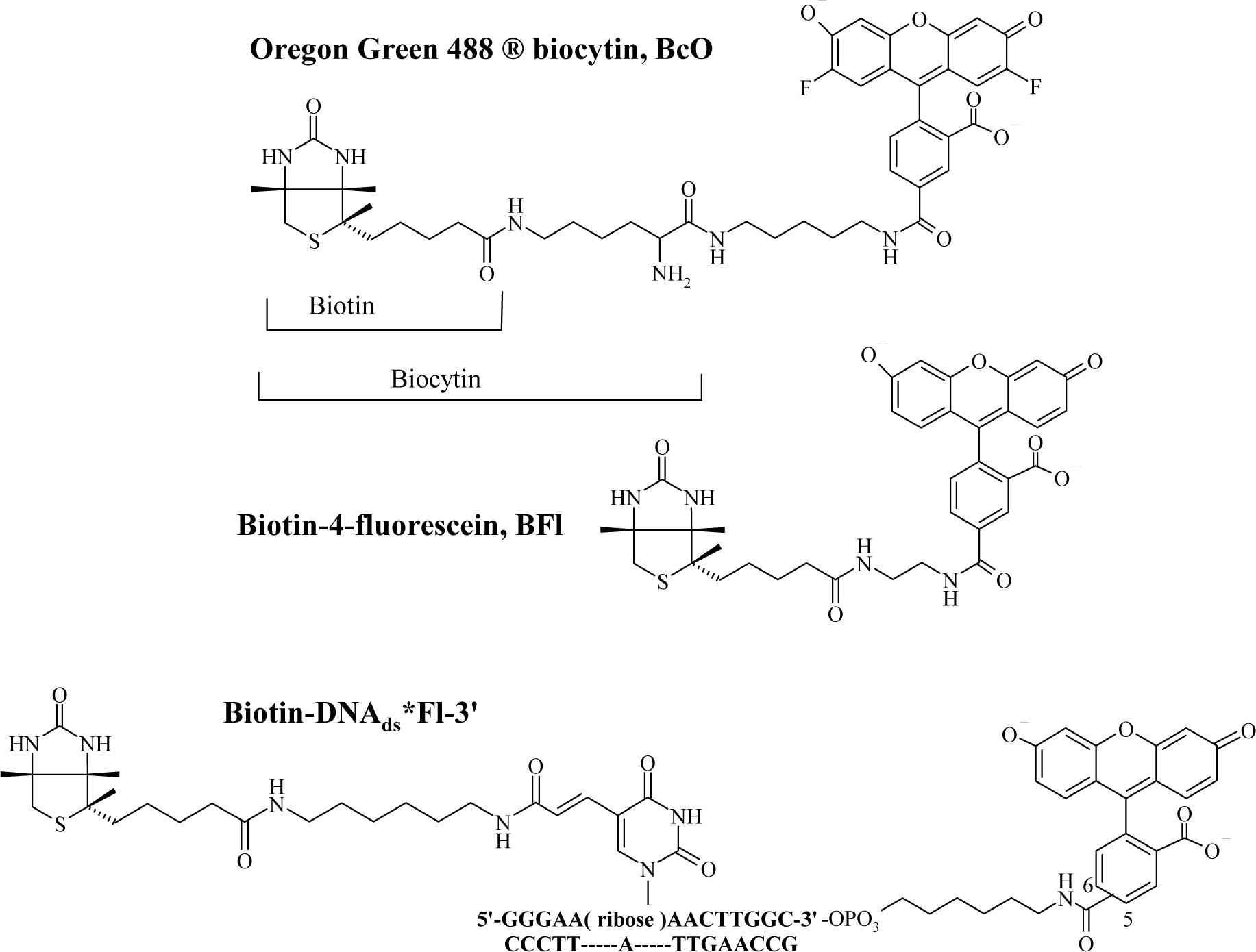
Dye-labeled B_7_ probes: 1) Biotin-4-fluorescein (BFl) contains a shorter spacer of 10 non-hydrogen atoms between the bicyclic ring and the dye structure. 2) Oregon green 488 ® Biocytin (BcO) has a spacer of 20 non-hydrogen atoms between the bicyclic ring and the fluorescent dye. Biocytin (Bc) is an amide formed with B_7_ and L-lysine. 3) Biotin-DNA_ds_*Fl-3’, where B_7_ was attached to a 14-mer DNA duplex labeled with fluorescein (Fl) at the 3’ end with 16 non-hydrogen atoms between the bicyclic ring and the thymine cyclic base. Unlabeled B_7_ was used to find the reaction rate of the final binding site in AV and compare it with the reaction rates of the initial binding site to assessment possible cooperativity.

We also studied the effect of AV glycosylation by removing, enzymatically, the carbohydrate motif to compare the respective association rates with those of the untreated AV, SAV and analogous probes in other studies.^20,33–36^ We show that the binding polynomial distribution (Z) of the dye-labeled B_7_ complexes to track bound tetrameric species that appeared after SF mixing at pseudo first order conditions corresponding to high excess protein levels. We make a distinction of the AV and SAV complexes using a simple filling model AB_n_ where A is either AV or SAV, and “n” is the total available number of binding sites occupied by the dye-labeled B_7_ probes and not the Hill number associated with cooperative binding.

For the second methodology, using a relaxation kinetics approach, the association reactions of unlabeled B_7_ was monitored in SF instrumentation by tracking the absorbance changes of an AV-HABA complex as B_7_ replaces bound HABA.^38^ The presence of ligand stabilizes the avidin tetramer. AV-HABA relaxation experiments were used to determine if stabilizing the tetramer affects the association rate constants and cooperativity.

Global fitting of the kinetic traces and reported calorimetry values allowed us to test reaction models and discriminate the most probable reaction mechanism, as carried out by some of us in previous studies.^39–42^ Consequently, the respective activation energies calculated by Arrhenius plots of association rates allowed the acquisition of the forward thermodynamic parameters toward the transition state: enthalpy (E_a_^forward^ or ΔH^‡, forward^), entropy (ΔS^‡, forward^) and Gibbs energy (ΔG^‡, forward^) of AV and SAV activated complexes. The forward thermodynamic data is in excellent agreement with the backwards thermodynamic values calculated with the dissociation rate constants (k_off_) reported by N. M. Green in his seminal work.^32^ Additionally, we explain the nature of the second dissociation phase first observed and correctly neglected by Green as a bimolecular “displacement” rate constant 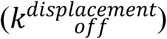, in addition to the detection of the documented unimolecular “replacement” rate constant 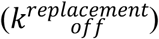^26,32^ which is used to establish the well-known dissociation constant, K_D_, as the most stable complex in nature.

Furthermore, we studied the changes in fluorescence lifetime (τ), quantum yield (QY), dynamic quantum yield (Φ), dye emitting fraction (1-S) and steady state anisotropy (r_ss_) of the fluorescent probes before and after complex formation. These spectroscopic properties give indications of the chemical environment surrounding the B_7_ binding pocket in AV and SAV and have important relevance in fluorescence assay detection limits as the signal to noise ratio can be improved by careful choosing linker length and fluorescent probe.

## 2. EXPERIMENTAL PROCEDURES

### 2.1.1. Materials and solution conditions

Oregon green ® 488 Biocytin (lot 40300A, Figure 1) was from Invitrogen (Eugene, OR). Avidin (CAS 1405-69-2, lot 608540) was purchased from Calbiochem (La Jolla, CA). HABA or 2-(4’-hydroxyazobenzene)-benzoic acid (CAS 1634-82-8, lot 52F-0073), streptavidin (CAS 9013-20-1), endoglycosidase H (CAS 37278-88-9) and d-biotin (CAS 58-85-5, lot 13F-3199) were all purchased from Sigma Aldrich (St. Louis, MO). Biotin-4-fluorescein (lot 31005, Figure 1) was purchased from Biotium, Inc. (Hayward, Ca). The 3’ end labeled fluorescein top strand with a modified biotinylated d-thymine at position 6 in the following sequence: 5’-GGGAA(biotin-dT)AACTTGGC*Fl-3’ (Figure 1) and the respective complement (5’-GCCAAGTTATTCCC-3’) were made by Tri-Link Biotechnologies, Inc. (San Diego, CA), and were both HPLC and PAGE purified. The sequences retain the G/C (base pairs) ends and fluorescein identical to those characterized extensively in our previous studies. ^39,41,43^ The biotinylated 14mer duplex (biotin-DNA_ds_*Fl) was formed with 5-10X excess complement and incubated for at least 20 min.

### 2.1.2. The protein and active site concentrations of AV and SAV

were determined with the HABA colorimetric assay of Green^37^ for which absorbance measurements, with total protein at 280 nm (1.54 = 1 mg/ml) and HABA at 500 nm (35.5 mMε bound, 0.48 mMε unbound) were made with a Cary 300 Bio UV-Vis spectrophotometer (Varian Inc., Palo Alto, CA). The occupancy of the dye-labeled probes on the AV and SAV tetramer (“p”) was obtained with the expansion version of the normalized partition function, Z = (p + q + x)^4^. In considering the totality of binding sites in the AV and SAV tetramer, let “p” denote the fraction of total sites occupied by B_7_ ligands (or HABA), “q” the fraction that are unoccupied and are available for binding, and “x” the fraction that are unavailable. The normalized partition function that describes the mole fractions of the various possible AV and SAV tetrameric species is given by Z = (p + q + x)^4^; where “x”, from the HABA assay for AV, was found to be 0.185 (or 18.5%), and q = 1 − p − x. Knowing the total concentration of binding sites from UV protein absorbance and Green’s methodology,^37^ and determining “x”, gives one the maximum value of “p” that will be reached in reacting tetramers with a B_7_ analog. Expansion of Z gives the mole fractions of the various species in solution, and in decreasing order in terms of probe occupancy, are: p^4^ + 4p^3^q + 4p^3^x + 6p^2^q^2^ + 6p^2^x^2^ + 12p^2^qx + 4pq^3^ + 4px^3^ + 12pq^2^x + 12pqx^2^ + q^4^ + x^4^ + 4q^3^x + 4qx^3^ + 6q^2^x^2^ which totals 1. This development assumes completely random occupancy of probe and inactive sites characterized by “x”. The species containing one bound probe have “p” raised to the first power; those with two bound probes have “p” raised to the second power, and so on.

All of the following protein concentrations are presented on a binding site basis: The HABA association reactions for AV were measured at 23 ± 0.1 °C with a concentration of 87 μM HABA and 7.7 μM AV. The AV-HABA relaxation reactions were conducted with a preformed AV-HABA complex made up of 200 μM HABA and 10 μM AV, flowed against varying amounts of biotin from 100 μM up to 4000 μM for a [HABA]/[B] ratio that ranged from 0.05 to 2.

### 2.1.3. The association stopped-flow reactions

were carried out in a buffered solution of 10 mM Tris-HCl, 100 mM KCl, 2.5 mM MgCl_2_ and 1 mM CaCl_2_ at pH 8 and only AV-BcO reactions included pH 9 and 10. The concentrations, after mixing, were of 20 nM of dye-labeled biotin probe and 260 nM, 520 nM or 1040 nM of AV; and for SAV of 200 nM, 300 nM, 400 nM or 800 nM at temperatures of 10, 15, 20 and 25 °C. The deglycosylation of AV (for comparative association reactions) was carried out with endoglycosidase H^44^ with the standard protocol provided, both with and without incubation of a denaturant solution (2% SDS and 1M 2-mercaptoethanol).

### 2.1.4. The dissociation reactions of dye-labeled biotin complexes

were carried out with AB_1_ and AB_4_ preformed complexes. The AB_1_ complexes were created with 800 nM SAV and 40 nM of BcO or BFl (∼11% mole fraction) and for the AB_4_ complexes with 40 nM SAV and 40 nM of BcO or BFl. The AB_1_ complexes, at 20 ± 0.1 °C, were challenged with several concentrations of unlabeled B_7_ of 1500 nM, 1750 nM, 2000 nM and 2500 nM, of which 760 nM filled the remaining binding sites not occupied with the fluorescent probe and at 27 ± 0.1 °C, the total challenging B_7_ concentrations were of 1300 nM, 1500 nM, 1750 nM, 2000 nM and 3000 nM. The AB_4_ complexes, at 20 ± 0.1 °C, were challenged with biotin concentrations of 400 nM and 1600 nM.

### 2.1.5. The lifetimes (τ), steady state anisotropies (r_ss_), time-resolved anisotropies (r_t_) and quantum yields (QY) of the complexes

(at 20 ± 0.1 °C and pH 8) were collected with a dye-labeled B_7_ probe concentration of 20-40 nM and 1040-2080 nM of either protein (AV or SAV) to ensure that only one binding site in the tetramer was filled with a ligand (AB_1_ filling model).

### 2.2. Methodologies

#### 2.2.1. Steady-state anisotropy (r_ss_)

measurements were collected using the Giblin-Parkhurst modification of the Wampler-Desa method as described previously.^45^ The fluorescence signal was detected in a model A-1010 Alphascan fluorimeter (Photon Technologies, Inc., Birmingham, NJ) equipped with an R928 PMT (Hamamatsu, Bridgewater, NJ). The excitation was provided by an Ar^+^ ion laser (Coherent Innova 70-4 Argon, Santa Clara, CA) at 488 nm and 5-10 mW of power incident on the sample. A photoelastic modulator (PEM-80; HINDS International, Inc., Portland, OR) was placed between the laser source and the sample compartment with a retardation level of 1.22π, and with the PEM stress axis orientated 45° with respect to the E vector of the laser beam. Two signals were acquired with the PEM alternating between “on” and “off” positions for 10 seconds and the data fitted to a least squared straight line to minimize noise and at least six of these independent measurements were averaged to acquire the r_ss_ values. The fluorimeter G factor was determined using a film polarizer and analyzer with an excitation at 488 nm provided by a xenon arc lamp (model A1010, Photon Technologies Inc, Princeton, NJ). The dissociation reactions of dye-labeled B_7_ and protein complexes were monitored by fluorescence changes and were also collected in the fluorimeter described above.

#### 2.2.2. Fluorescence lifetimes (τ) and time-dependent anisotropy decays (r_t_)

were collected in a FluoTime100 fluorescence spectrometer (PicoQuant, GmbH, Berlin, Germany) with the excitation light source provided by a picosecond pulsed diode laser (PicoQuant, GmbH, Berlin, Germany) at 470 nm and 20 MHz. The emission was collected at 520 nm through a non-fluorescing 520 nm interference filter (Oriel Corp., Stratford, CT) followed by a liquid filter of 1cm path length containing 24 mM acetate buffered dichromate at pH 4, placed between the sample and detector to eliminate traces of excitation light.^39^ The fluorescence decays were fit by a nonlinear least-squares minimization based on the Marquardt algorithm embedded in the Fluofit software (PicoQuant GmbH). Twenty-eight decays were collected per sample; they were grouped in four sets, each set consisted of seven sample decays and one Instrument Response Function, IRF, for deconvolution proposes. The sets were globally fitted to mono- or bi-exponential decay models that were discriminated using the statistical parameter χ^2^. The r_t_ data was acquired with the fluorimeter described above equipped with a polarizer and an analyzer to acquire the parallel VV(t) and perpendicular VH(t) decays. The PicoQuant G factor was calculated according to: G=∫HV(t)dt/∫HH(t)dt, where HV(t) and HH(t) were the decays collected with the emission polarizer selecting vertical and horizontal E-vector passing orientations, respectively; and the excitation polarizer set for horizontally polarized excitation.

#### 2.2.3. Quantum yields (QY)

were obtained by using a reference fluorophore of known quantum yield and were calculated according to Parker and Rees,^46,47^ where the reference dye was fluorescein in 0.1N sodium hydroxide solution.^38^ The emission fluorescence scans were collected from 480 nm to 700 nm with the excitation light set at 460 nm provided by the xenon arc lamp described above. These measurements were made on the AB_1_ complexes at high protein concentration.

#### 2.2.4. The intrinsic lifetime (τ°), dynamic quantum yield (Φ) and the fraction of non-statically quenched molecules

*(1-S)* calculations have been described elsewhere^43^ and were acquired for the AB_1_ complexes. The HABA association reaction for AV was carried out under pseudo-first order conditions on a micro absorbance SF instrument^48^ equipped with a xenon arc lamp (described above) and a monochromator (model 82-410, Jarrel-Ash, Waltham, Mass.) set at 500 nm.

#### 2.2.5. The relaxation kinetics of unlabeled biotin reacting with the AV-HABA complexes

were prepared at concentrations in which HABA occupies all sites (AV-HABA_4_). Biotin replaces HABA relative to the *k*_*off*_ of the dye as shown for the first step (Eqn. 1) and then repeated for all sites. Having greater affinity, B_7_ occupies all sites at the end of the reaction and the measured *k*_*on*_ is related to the affinities of the ligand bound (R-state) protein.

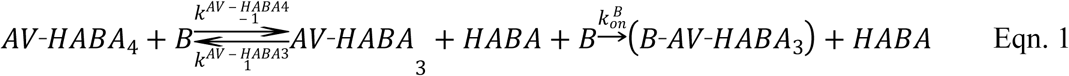

The reaction is monitored by the HABA absorbance changes at 500 nm as it is replaced by unlabeled B_7_; yielding the relaxation constant of the reaction (R, Eqn. 2) which contains information of the biotin association rate constant of the R-state open binding site, 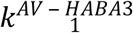 to form a full saturated complex (AV-HABA_4_) and the dissociation rate of that full complex, 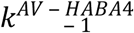, to yield a complex with three HABA molecules (AV-HABA_3_). In subsequent steps, biotin replaces HABA as the ligand but the release of HABA creating an unoccupied site remains the same treating the reaction on a per monomer basis. The experiment was designed to acquire the pseudo first order association rate constant of B_7_ binding 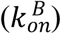 to the R-state binding site in a complex occupied by HABA molecules (AV-HABA_4_).

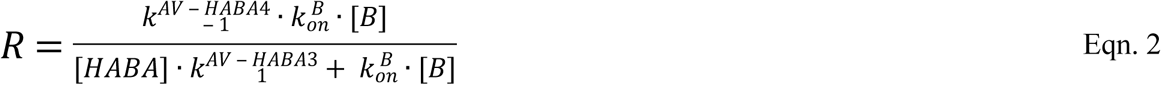

The reciprocal of the relaxation constant (1/R) is plotted vs. the [HABA]/[B] concentration ratio (Eqn. 3) allowing to calculate: 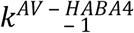 and 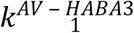 by solving for the intercept 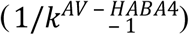 and the respective slope: 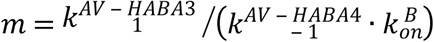. The exponential decays were analyzed by the method of Foss.^49^ There was no departure from simple first order decay in the relaxation, justifying the use of the following simple model and equations.

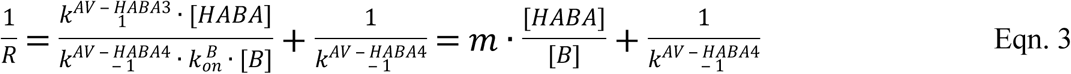

#### 2.2.6. The association reactions of dye-labeled biotin and AV (or SAV)

were collected with a SF instrument, described previously.^50,51^ The fluorescence signal was collected through a 520 nm interference filter (Oriel Corp., Stratford, CT) with a detector time constant and SF dead time of 1 μs and 1 ms, respectively. The excitation light was provided by the Coherent Ar^+^ ion laser (described above) at 488 nm with 15-10 mW of incident power on the reaction cuvette. The laser source was followed by the photo-elastic modulator described above with the axis oriented 45° with respect to the electric vector of the incident light and with the half-wave modulation (50 kHz) set for 488 nm excitation. The demodulation circuitry following the photomultiplier provided a DC(t) and a rectified AC(t) which were converted to digital data by a high-speed digitizer (PCI-5122) from National Instruments (Austin, TX) with 14-bit resolution and 100 MHz bandwidth, through channels 0 and 1. The data acquisition was controlled by LabVIEW(tm) (Vr 8) software at a collection rate of 6120 data points/second and stored in spreadsheets. The AC(t) and DC(t) data were baseline corrected before obtaining the signal ratio (Eqn. 4) as a function of time (ρ_t_).

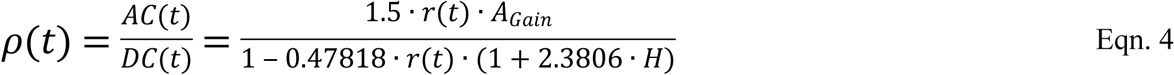

The constant A_Gain_ is the instrumental amplitude gain and was evaluated by solving ρ(t) using the known steady state anisotropy (r_ss_) of the complexes which is equivalent to the r(t) at t= ∞; and H, obtained from the equivalent grating factor (G) for the filters and photo multiplied tubes in the SF. For the probes used in here G was 0.82 and H = (1-G)/(1+G) = 0.099. Knowing A_gain_ and H, the AC(t) and DC(t) signals can be solved for *r*(*t*) and *F*(*t*) (Eqn. 4 and 5) and the normalized fluorescence, 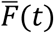, and corrected fluorescence anisotropy, 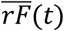,^52^ were obtained when 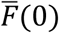 and 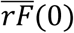 were scaled to 1 at t=0.

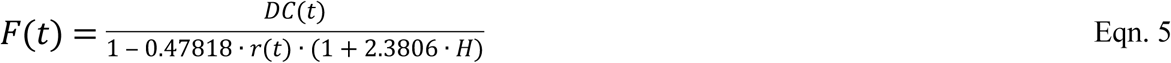

The 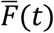 (Eqn. 6) is equivalent to (I_∥_) + 2 (I_⊥_) and proportional to quantum yield (QY_i_), molar absorptivity (ε_i_) and to the formation or disappearance of the emitting species X_i_(t); and 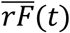 including the steady state anisotropies (r_ss_) of each fluorescent species (Eqn. 7).^52^

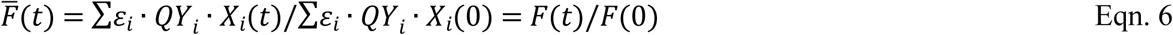

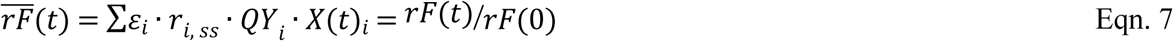

#### 2.2.7. The biotin association reaction model for AV and SAV

was discriminated by the squared residuals of the observed and calculated association traces of both fluorescence and anisotropy fluorescence signals, 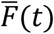 and 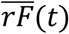, respectively. For the labeled biotin probes: BFl and BcO probes, the association reactions were very well described by the simplest possible model (Eqn. 8) with single association rate constants (k_on_).

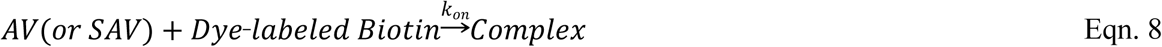

In the case of the biotin-DNA_ds_*Fl, the association reaction model was complemented by second k_on_ which resulted in a system of two parallel reactions (Eqn. 9). In both cases, the backward reaction is not significant during the 5-8 sec required for the biotin association binding.

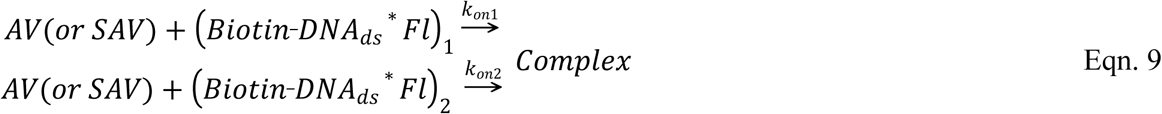

#### 2.2.8. The dissociation reactions of the complexes

were followed by fluorescence changes, 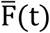, in the fluorimeter and laser setup described above and tuned to 488 nm under discontinuous excitation to prevent photobleaching distortion. The signal was best fitted to the following dissociation model (Eqn. 10), in which the dye labeled complex dissociates into the labeled B_7_ probe (BFl or BcO) and the respective protein (AV or SAV).

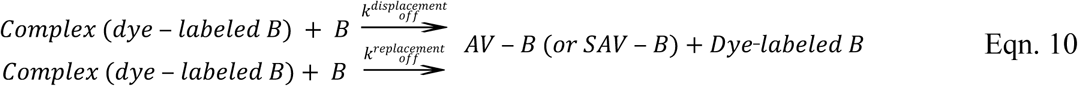

#### 2.2.9. Time-resolved anisotropy

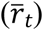 was calculated according to Eqn. 11 where the pre-exponential “*f* “corresponds to the slow phase that derives from the lifetime of the global motion (τ_G_)^53^ which was fitted within a range of expected correlation time for the complex size;^54^ consequently, facilitating resolution of the fast correlation lifetime (τ_p_) and the corresponding pre-exponential (1-f).

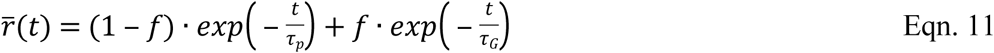

The *f* parameter was constrained to the observed r_ss_ (Eqn. 12) where 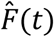 (Eqn. 13) is normalized (*α*_1_ + *α*_2_ = 1) and derived from the observed fluorescence decays of the complex.^55^

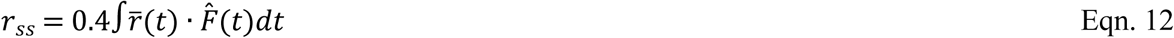

Where

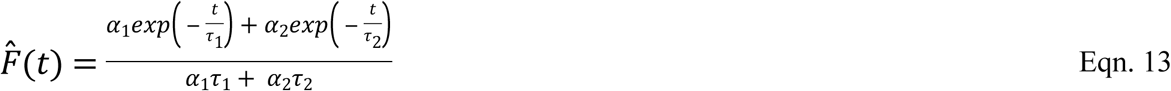

In a simple model, the transition moment is assumed to wobble within a cone of semi-apical angle Ω,^56^ where the cone axis is normal to the surface of a sphere that corresponds to the macromolecule. The angle Ω is calculated from Eqn. 14.

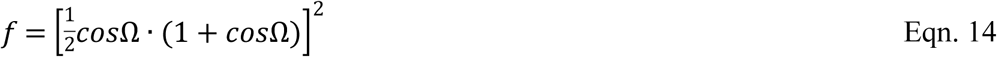

## 3. RESULTS AND DISCUSSION

### 3.1 Active avidin binding sites

Avidin and streptavidin are tetramers in solution. If the binding of ligand is positively cooperative, differences in *k*_*on*_ for initial (T-state) and final (R-state) binding steps could be significant and therefore comparison of initial binding by unliganded avidin and final binding by liganded avidin is necessary. Measurement of the initial binding rate requires ligand free avidin but endogenous ligand could potentially interfere. In fact, AV preparations often present about 20% of the inactive sites for the binding of any B_7_ analogs, either because they contain endogenous B_7_,^37^ or perhaps existence of damaged binding sites in some of them, *e.g*., tryptophan oxidation.^57^ To acquire accurate k_on_ values, the actual available binding site concentration for each sample was measured by HABA colorimetric assays in relation with absorbance at 280 nm. Accordingly, the percentage of available active sites of AV and SAV were 81.5 ± 0.5 % and 94.0 ± 1.0 % with respect to total protein, respectively, which were in excellent agreement with the 82 % and 95 % reported by the commercial source (Sigma Aldrich and CalBiochem). The SF apparatus provided rapid thorough mixing of the probes with AV and SAV allowing measurement of the full reaction. The issue of rapid mixing vs. more conventional titrations was treated previously.^48^ In the SF association measurements, the dye-labeled B_7_ probes were sub-stoichiometric to determine the initial (T-state) binding rates (e.g. 20 nM of BFl, BcO and biotin-DNA_ds_*Fl vs. 260 nM, 520 nM and 1040 nM in binding sites basis). Limiting the ligand also reduces several potential measurement artifacts including FRET self-transfer, and contact interference including probe fluorescence quenching by contact interactions^58^ in the AB_2_, AB_3_ or AB_4_ complexes; especially for the BcO which has a longer linker.^59^ Using the binding polynomial for the 20 nM probe after mixing, and for the intermediate AV concentration, 638 nM in total sites, 520 nM in available sites, the mole fraction of species with a single bound probe is 0.114, that with two bound is 0.0055, with three bound is 0.00012, so at most, only 4.6 % of the molecules with bound AV contain two probes; for 1040 nM available sites, the value drops to 2.3 %. With limited occupancy, the association reactions acquired the dye-labeled B_7_ probes reflect the binding to the first binding site (T-state) in the tetramer for the SF experiments. Unlabeled B_7_ relaxation kinetic experiment was designed to observe the binding at the final site (R-state), as discussed below.

### 3.2. Association rate constants (k_on_) of biotin binding to apo-avidin

#### 3.2.1. Dye-labeled biotin association rate constants by stopped-flow methodology

The fluorescence 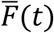 and corrected anisotropy association binding traces, 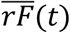, properly monitored the association reactions, as they yielded equivalent k_on_ values (Table 1) and presented the best optimal fit residuals (Figure 2). In contrast to the anisotropy signal, 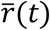, that lagged behind 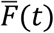 and 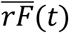 owing to changes in the quantum yield (QY) of the involved fluorescence species.^52^ These three types of association binding traces were acquired with discontinuous excitation that circumvented photobleaching (Figure 3) allowing the detection of all non-photobleaching rate constants. Consequently, the k_on_ values of AV showed linear concentration dependence (Figures 4) and strong temperature dependence when using the BcO (Figure 5) and BFl (Table 2) probes. Notably, a reduction in the k_on_ of ∼10% was observed with each pH unit increment (from 8 to 10) which may derive from titration of the hydrogen bonding of asparagine and tyrosine in the binding pocket.^29^

**Table 1.** Comparison of the association rate constants (k_on_) obtained by fluorescence change, 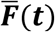, and corrected fluorescence anisotropy, 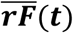, of BcO (20 nM) binding to AV at several temperatures, protein concentrations and pH 8.^a^

| Complex<sup>a</sup> | 25 °C<br>($\times 10^{-6} \text{M}^{-1} \text{s}^{-1}$) | 20 °C<br>($\times 10^{-6} \text{M}^{-1} \text{s}^{-1}$) | 15 °C<br>($\times 10^{-6} \text{M}^{-1} \text{s}^{-1}$) | 10 °C<br>($\times 10^{-6} \text{M}^{-1} \text{s}^{-1}$) | $E_a^{\text{Forward}}$<br>(kcal/mol) | $\Delta S^\ddagger$<br>(cal/°K·mol) | $\Delta G^\ddagger$<br>(kcal/mol) |
| --- | --- | --- | --- | --- | --- | --- | --- |
| $k_{on}$, AV-BcO | $\bar{F}(t), \times 10^{-6} \text{M}^{-1} \text{s}^{-1}$ | | | $\overline{rF}(t), \times 10^{-6} \text{M}^{-1} \text{s}^{-1}$ | | | |
| Temp. | 260 nM | 520 nM | 1040 nM | 260 nM | 520 nM | 1040 nM |  |
| 25 °C | 9.5 ± 0.1 | 9.5 ± 0.3 | 9.7 ± 0.3 | 9.5 ± 0.1 | 9.5 ± 0.1 | 9.4 ± 0.2 |  |
| 20 °C | 5.7 ± 0.1 | 5.9 ± 0.3 | 6.0 ± 0.1 | 5.9 ± 0.1 | 5.8 ± 0.1 | 6.1 ± 0.1 |  |
| 15 °C | 4.1 ± 0.1 | 3.95 ± 0.1 | 3.9 ± 0.2 | 4.0 ± 0.1 | 3.82 ± 0.1 | 4.0 ± 0.2 |  |
| 10 °C | 2.4 ± 0.1 | 2.6 ± 0.2 | 2.5 ± 0.2 | 2.7 ± 0.1 | 2.7 ± 0.1 | 2.7 ± 0.1 |  |

<sup>a</sup>The $\bar{F}(t)$ and $\overline{rF}(t)$ signals were equivalent as they tracked in the errors the association process of dye-labeled B<sub>7</sub> binding to the proteins under pseudo first order conditions.

**Table 2.** Association rate constants (k_on_) and thermodynamic values of the dye-labeled biotin binding to AV and SAV.

|  |  |  |  |  |  |  |  |
| --- | --- | --- | --- | --- | --- | --- | --- |
| **AV-BFI** | 18.5 ± 2.0 | 10.7 ± 1.6 | 7.60 ± 2.0 | 4.86 ± 0.6 | 14.6 ± 0.6 | 34.9 ± 2.0 | 4.2 ± 0.5 |
| **AV-BcO (pH 8)<sup>b</sup>** | 9.5 ± 0.4 | 5.9 ± 0.4 | 3.9 ± 0.3 | 2.6 ± 0.3 | 14.4 ± 0.6 | 33.0 ± 2.5 | 4.6 ± 0.3 |
| **AV-BcO (pH 9)** | 7.9 ± 0.3 <sup>c</sup> | 5.4 ± 0.2 | 3.4 ± 0.1 | 2.3 ± 0.1 | 13.9 ± 0.7 | 30.9 ± 2.5 | 4.7 ± 0.1 |
| **AV-BcO (pH 10)** | 7.2 ± 0.3 <sup>c</sup> | 4.8 ± 0.2 | 3.0 ± 0.1 | 1.9 ± 0.1 | 14.9 ± 0.7 | 34.1 ± 1.3 | 4.7 ± 0.3 |
| **1) AV-Biotin-DNAd<sub>s</sub>\*FI** | 15.7 ± 1.0<br>(30.5 %) d | 11.2 ± 0.8<br>(68.9 %) | 7.4 ± 0.7<br>(87.5 %) | 4.23 ± 0.5<br>(95.0 %) | 14.6 ± 0.8 | 34.8 ± 3.0 | 4.2 ± 0.2 |
| **2) AV-Biotin-DNAd<sub>s</sub>\*FI** | 1.4 ± 0.1<br>(69.5 %) | 0.98 ± 0.05<br>(31.1 %) | 0.7 ± 0.03<br>(12.5 %) | 0.52 ± 0.01<br>(0.05 %) | 14.2 ± 0.5 | 29.0 ± 0.5 | 5.6 ± 0.6 |
| **Average** | NA | NA | NA | NA | 14.4 ± 0.2 | 32.8 ± 1.2 | 4.7 ± 0.3 |
| **SAV-BFI** | 74.7 ± 2.0 | 58.6 ± 1.6 | 53.1 ± 2.0 | 45.4 ± 1.0 | 5.3 ± 0.3 | 6.6 ± 0.5 | 3.3 ± 0.6 |
| **SAV-BcO** | 30.3 ± 2.0 | 24.0 ± 1.6 | 20.0 ± 1.2 | 17.3 ± 0.6 | 6.2 ± 0.4 | 7.8 ± 0.4 | 3.9 ± 0.4 |
| **1) SAV-Biotin-DNA<sub>ds</sub>\*FI** | 53.0 ± 1.0<br>(44.4 %) | 45.9 ± 0.8<br>(50 %) | 36.4 ± 0.7<br>(51 %) | 31.0 ± 0.5<br>(49 %) | 6.2 ± 0.4 | 8.8 ± 0.6 | 3.5 ± 0.5 |
| **2) SAV-Biotin-DNA<sub>ds</sub>\*FI** | 2.50 ± 0.1<br>(55.6 %) | 2.33 ± 0.1<br>(50 %) | 1.69 ± 0.1<br>(49 %) | 1.50 ± 0.05<br>(51 %) | 6.2 ± 0.5 | 3.1 ± 0.3 | 5.3 ± 0.6 |
| **Average** | NA | NA | NA | NA | 6.0 ± 0.2 | 6.6 ± 1.6 | 4.0 ± 0.6 |

**Figure 3.**
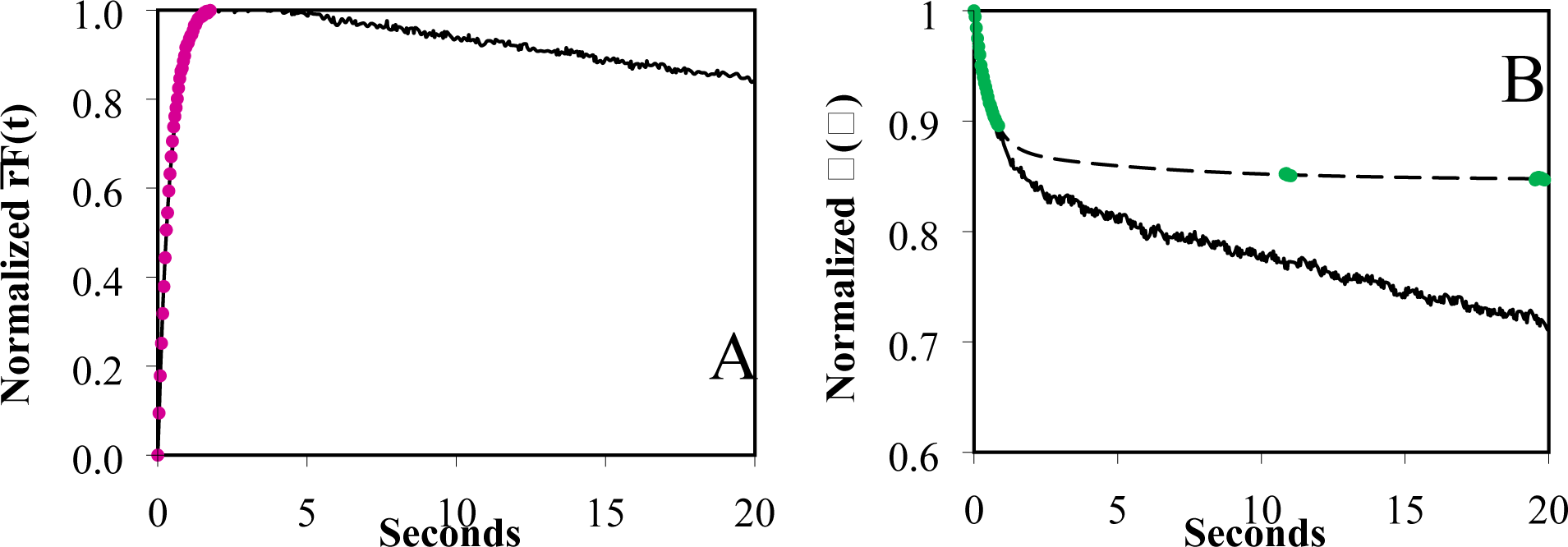
Photobleaching of BcO binding to AV at 15 °C: **A**) 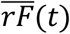 and **B**) 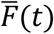 which corresponding normalization functions are: 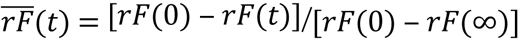 and 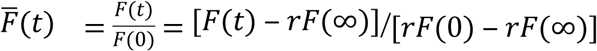, respectively. The photobleaching rate constant was elucidated by collecting the reaction with continuous (black) and discontinuous (dashed color) laser illumination, where in the latter case the beam was blocked during the times denoted by dashes and the sample was illuminated only during time intervals around 10 s. The slow photobleaching rate constant varied from 6 × 10^−3^ to 1 × 10^−2^ s^−1^, and was laser power dependent.

**Figure 4.**
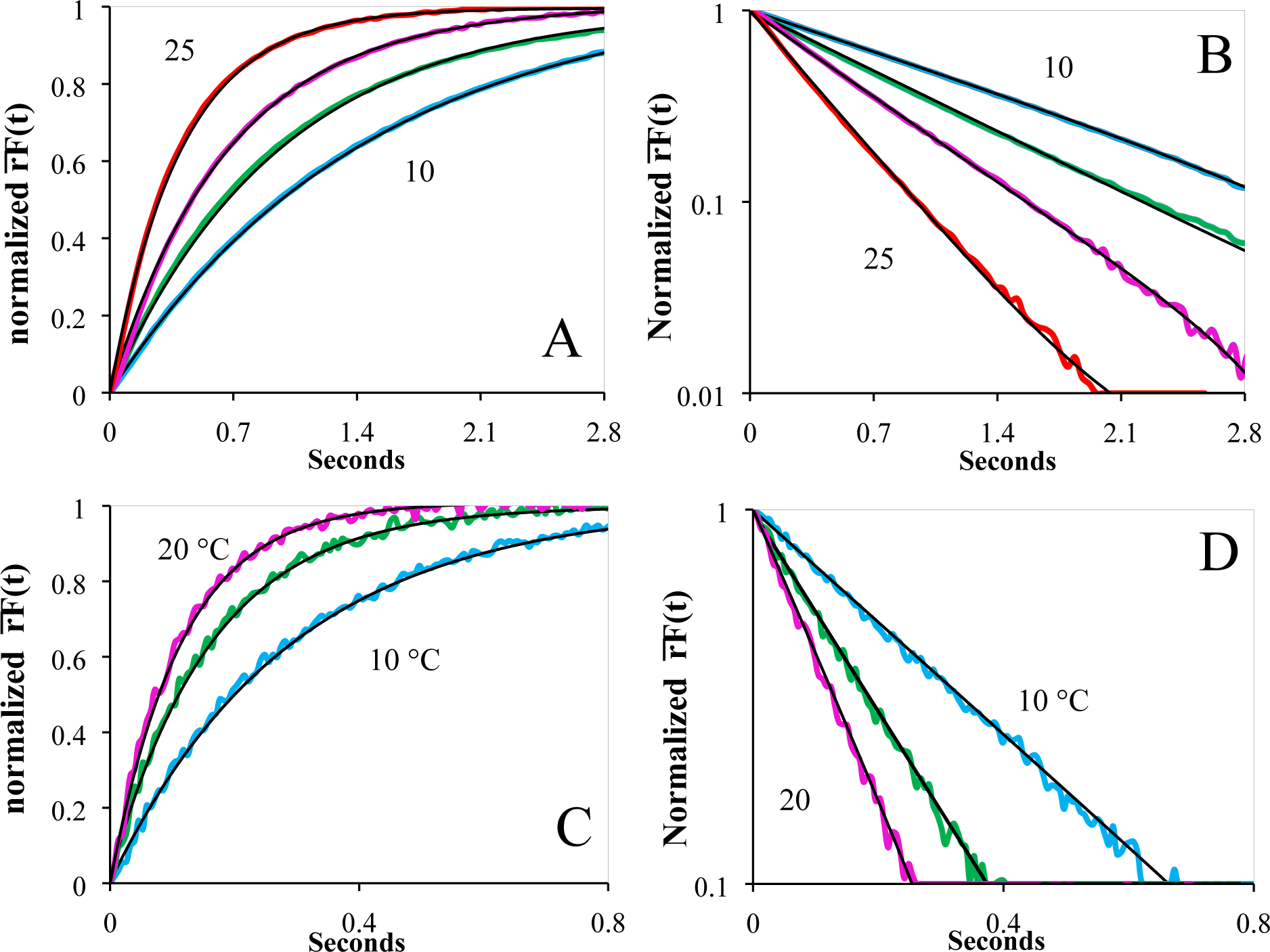
**A)** Normalized fluorescence anisotropy, 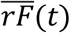, temperature dependence of BcO (20 nM) binding to AV (260 nM) at pH 8, normalized as 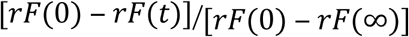. The observed (color) and fitted (black line) curves at 25 °C (top), 20 °C (upper middle) and 15 °C (lower middle) and 10 °C (bottom) had half-times of 280 ms, 452 ms, 695 ms and 1024 ms, respectively; and **B)** shows the corresponding semi-logarithmic plot of 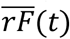. **C)** Normalized fluorescence anisotropy, 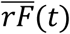, shows a temperature dependence of BcO (20 nM) binding to SAV (200 nM) at pH 8, normalized as above (4A). The observed (color) and fitted (black line) curves at 20 °C (top), 15 C (middle) and 10 °C (bottom) had half-times of 79.4 ms, 111 ms and 202 ms, respectively and **D)** shows the corresponding semi-logarithmic plot of the 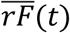.

**Figure 5.**
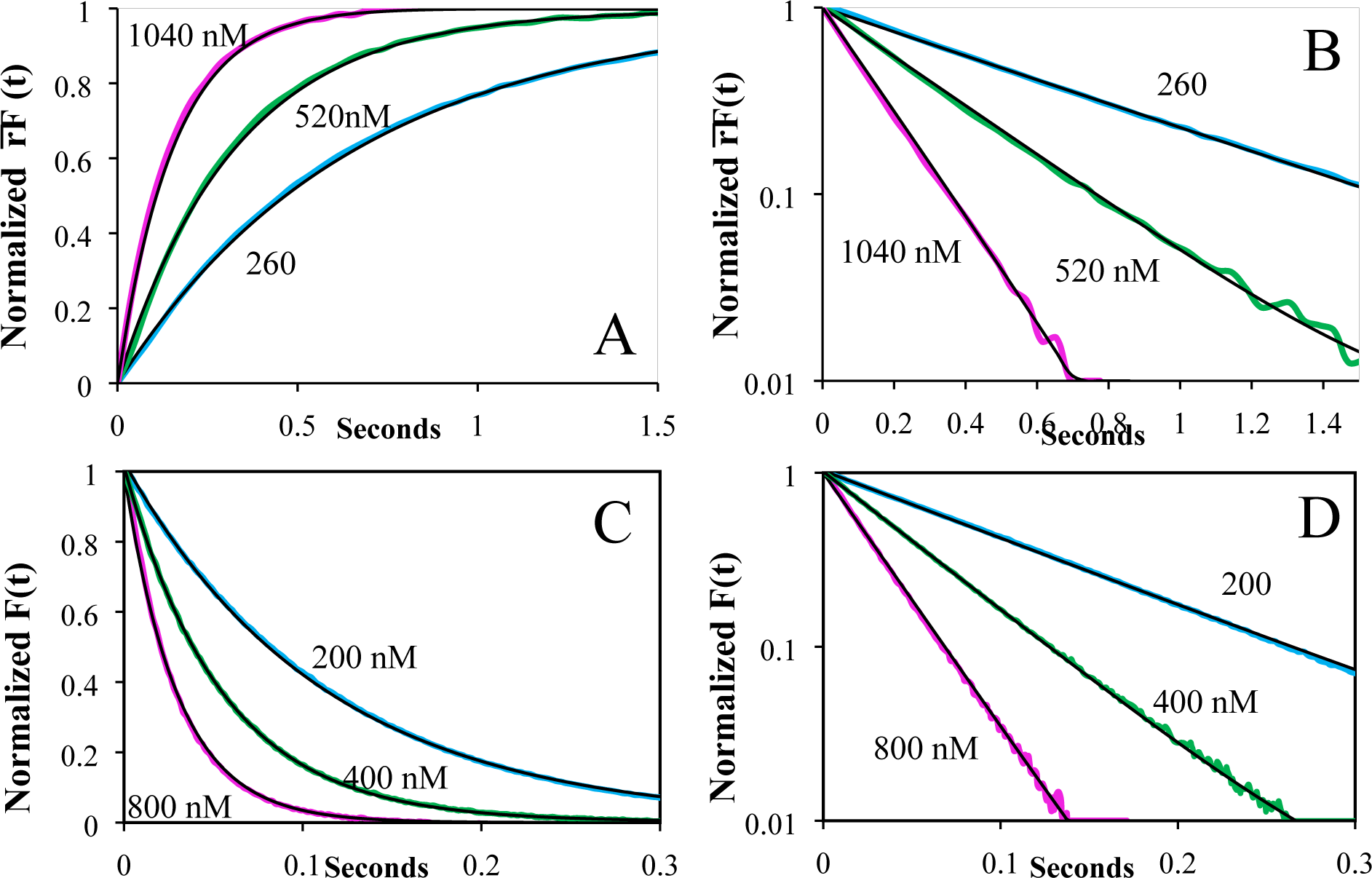
**A)** Corrected Fluorescence Anisotropy Concentration dependence of BcO (20 nM) binding to AV at pH 8 and 20 °C, normalized as 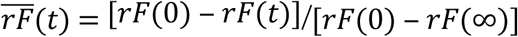. The observed (color) curves were obtained with an AV concentration of 1040 nM (top), 520 nM (middle) and 260 nM (bottom) and the fitted curves (black lines) had halftimes of 109 ms, 229 ms and 455 ms; respectively. **B)** Shows the corresponding semi-logarithmic plots of 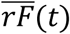. **C)** Normalized fluorescence change, 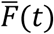, concentration dependence of BFl (20 nM) binding to SAV at pH 8 and 20 °C. Normalized as followed: 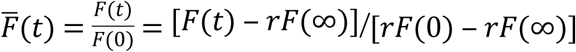. The observed (color) curves were acquired with SAV concentrations of 200 nM (red), 400 nM (purple) and 800 nM (orange) where the fitted curves (black lines) had halftimes of 77.3 ms, 37.7 ms and 20.6 ms; respectively. **D)** Shows the corresponding semi-logarithmic plot of 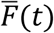

**Figure 2.**
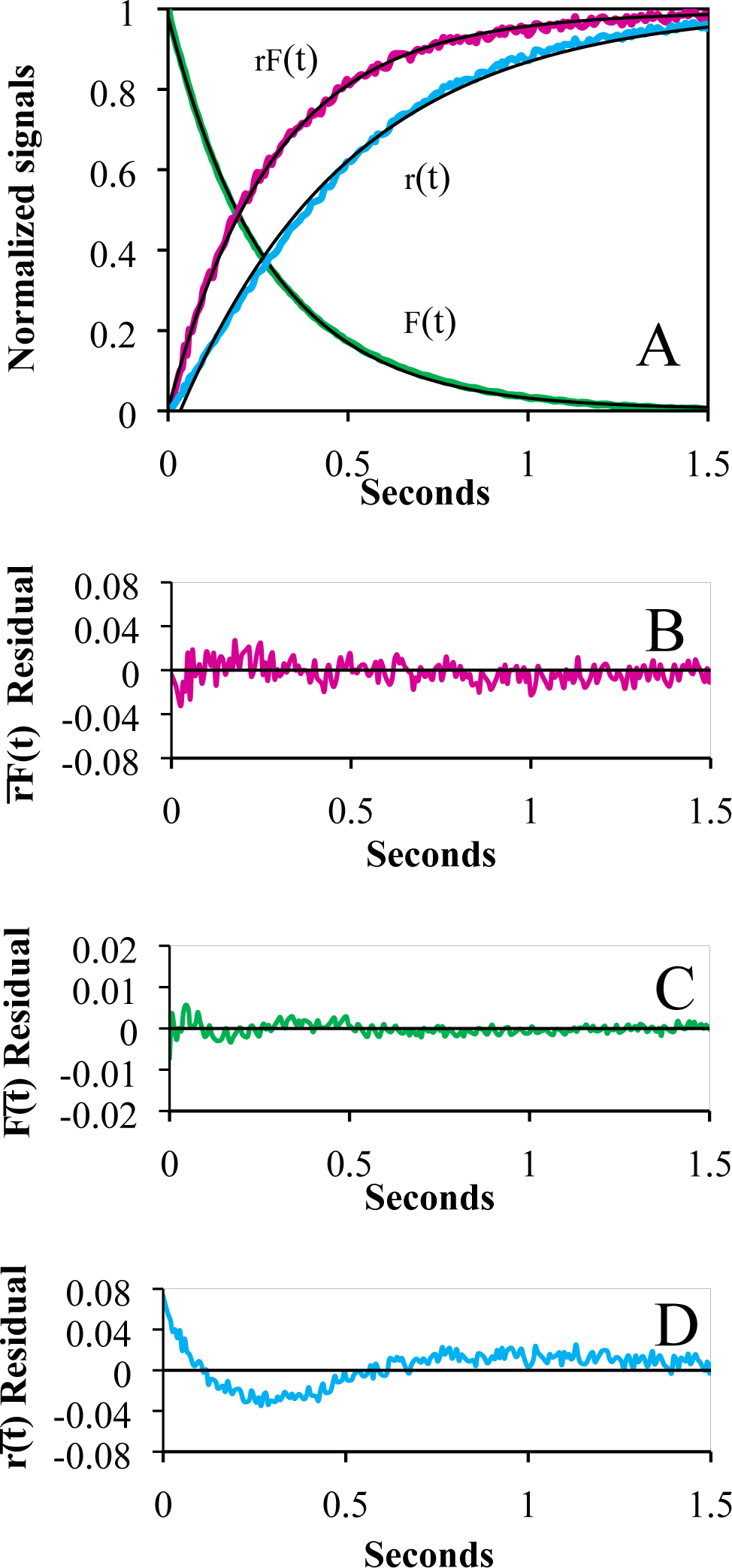
**A** Comparison of the fluorescence change, anisotropy and corrected fluorescence anisotropy signals, 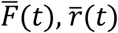 *and* 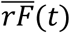, respectively; of BcO (20 nM) binding to SAV (200 nM) at 10 °C whose mono-exponential fits (black) resulted in k_on_ values of 1.73× 10^7^ M^−1^s^−1^, 1.72 × 10^7^ M^−1^s^−1^ and 1.04 × 10^7^ M^−1^s^−1^ with halftimes of 200.6 ms, 201.4 ms and 332.7 ms, respectively. The k_on_ of 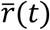 was 40 % slower than the other two and showed the worse residuals (**B-D**). The corresponding normalization signals are: 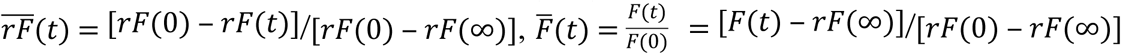 and 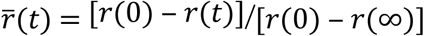.

#### 3.2.2. Unlabeled biotin association rate constants by relaxation kinetics methodology

The experiment consisted in challenging a pre-saturated AV-HABA complex with B_7_ (Figure 6) to measure the association rate of the final “relaxed” binding sites which yielded a 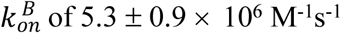 of 5.3 ± 0.9 × 10^6^ M^−1^s^−1^ (at pH 8 and 23 °C) which slightly slower than the 7.8 ± 0.4 × 10^6^ M^−1^s^−1^ acquired with BcO (Arrhenius plot, 23 °C and pH 8) indicating non-cooperativity (or slightly negative) for binding site association rates. The HABA dissociation rate constant of the AV-HABA_4_ complex was not rate limiting 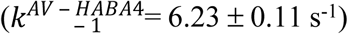 and the HABA association rate for the final site was 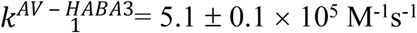 which results in a AV-HABA equilibrium constant of K_D_ ^AV-HABA^ = 12.2 ± 0.3 × 10^−6^ M similar to that reported by Green^57^ at pH 8 which support the quality of our relaxation kinetic experiment.

**Figure 6.**
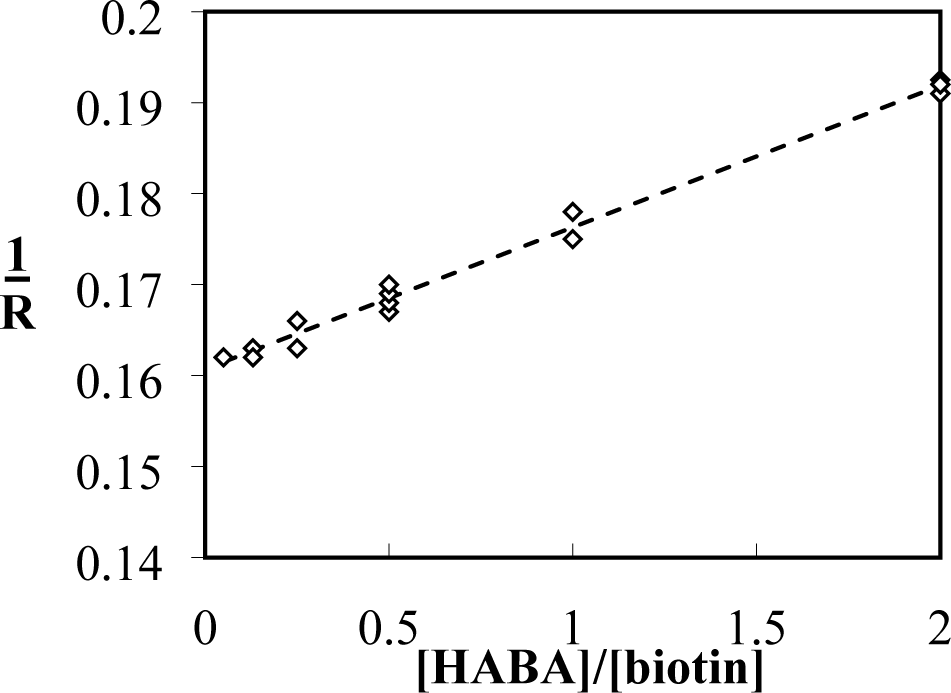
Relaxation kinetics of the AV-HABA complex and unlabeled biotin. The [HABA]/[B] is the concentration ratio of these two ligands and R is the relaxation rate in s^−1^ (Eqn. 3). The data points were fitted to a linear regression model yielding a slope and intercept of 0.0156 ± 0.0012 and 0.161 ± 0.005, respectively, resulting in a 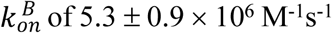 at 23 °C. The additional values required for this calculation were HABA association constant to the 4^th^ site when three HABA molecules are already bound: 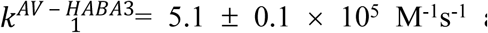 and the HABA dissociation of saturated AV-HABA_4_ complex: 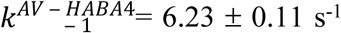. The corresponding K_D_ ^AV-HABA^ = 12.2 ± 0.3 × 10^−6^ M and is in excellent agreement with the 12 × 10^−6^ M reported by Green^57^ at pH 8.

#### 3.2.3. Non-cooperative biotin binding to avidin sites

The association reactions that used the fluorescent probes BFl and BcO monitored the 1^st^ available binding site, as they were carried out at pseudo first order, at very high protein concentration with low occupancy for AB_1_ filling model, as discussed above. In contrast, the relaxation kinetic methodology scrutinized the unlabeled B_7_ binding to the unoccupied site while the 3 remaining sites were filled with HABA, this process can be thought as the binding of B_7_ to the 4^th^ binding site. Therefore, the data obtained with dye-labeled B_7_ probes and unlabeled B_7_ should report the binding rates to the 1^st^ and 4^th^ sites. Since these two values only diverge by 32 % we believe that there is not significant cooperativity nor an intrinsic difference in any of the AV sites. If a protein has two forms, high (R) and low (T) affinity,^60^ the HABA bound ligand will hold the AV protein in the R-state. In the relaxation experiments, all the bound HABA get replaced by dye-labeled B_7_ (BFl or BcO) but all the sites are R-state meaning that there is not switching from T to R. This is the same as HbO_2_ flowed against CO, where O_2_ gets replaced by CO but not biphasic because no T-state is present.^60,61^ As B_7_ binding to AV and SAV is non-cooperative, the HABA replacement is a pseudo first order measure of the B_7_ association rate and as should be the same or close to the association rate of the dye-labeled B_7_ flowed against empty AV or SAV, and our values differed only by 32% for these two approaches.

#### 3.2.4 Comparisons with other AV-B kinetic studies

were carried out at the possible closest condition; thus, at 25 °C and pH 8, the BFl and BcO association rate constants, k_on_, were 3.8 and 7.4 × slower than the 7 × 10^7^ M^−1^s^−1^ reported by N. M. Green^32^ (at 25 °C and pH 5), respectively. However, a larger uncertainty is expected for the latter experiment because it was not carried out using rapid mixing techniques forcing the usage of very low (^14^carbon) B_7_ concentrations (picomolar range) to stop the reaction timely and quantify the un-reacted probe. Consequently, Green’s experiment was an extremely tedious task that was carried out, only once and at one temperature. On the other hand, a more recent association rate constant of 2.0 ± 0.3 × 10^6^ M^−1^s^−1^ was obtained in a Surface Plasmon Resonance study (SPR)^20^ at 20 °C and a pH 7.4 in HEPES buffer. This independent k_on_ value was ∼9 and ∼5 × slower than the ones acquired by us for BFl and BcO, respectively. Nevertheless, it has been acknowledged previously that the SPR results are, controversially, too low to be accurate.^20,36^ due to fixation to the chip of one of the reactants, generally AV or SAV.

#### 3.2.5 Effect of AV glycosylation on the biotin binding kinetics

AV has a glycan attached to asparagine 17 at each tetrameric subunit which is composed of four or five mannoses and three N-acetylglucosamines.^62^ These sugar modifications are typically removed to improve crystallization but the glycan effect on the association binding rate of B_7_ was previously unknown. Interestingly, after enzymatic removal of the carbohydrates, the k_on_ values of the de-glycosylated AV matched with those of natural glycosylated AV for the dye-labeled B_7_ probes: e.g., 3.7 ± 0.3 × 10^−6^ M^−1^s^−1^ vs. 3.9 ± 0.3 × 10^−6^ M^−1^s^−1^ of BcO binding to de-glycosylated AV and untreated AV at 15 °C, respectively. A previous study already suggested that the sugar chain is not required for B_7_ binding^62^ and now we confirm that AV glycosylation has no influence on the rate constants.

### 3.3. Association reaction of unlabeled and dye-labeled biotin binding to streptavidin

#### 3.3.1 Dye-labeled biotin association reactions to SAV presented temperature (Figure 4C and 4D) *and linear concentration dependence*

(Figure 5C and 5D). In comparison, at 25 °C, the k_on_ values of SAV were 4X and 3.2X faster than those observed for B_7_ binding to AV when reacting with BFl and BcO, respectively. However, the temperature dependence was weaker than that observe for AV which indicated a profound difference in the binding site properties of these two proteins. Thus, SAV should be a more robust system for purification applications as variations on the temperature incubation protocols has less negative significant effects in the yield.

#### 3.3.2 Comparisons with other SAV-B association kinetic studies

An independent SF study tracked the binding of unlabeled B_7_ by fluorescence quenching of the tryptophan (Trp) of SAV, yielding a k_on_ of 7.5 ± 0.6 × 10^7^ M^−1^s^−1^ (at 25 °C and pH 7)^36^ which was in excellent agreement with 7.5 ± 0.2 × 10^7^ M^−1^s^−1^ for the BFl probe (at 25 °C and pH 8). This finding strongly indicates that the attached dyes are innocuous and dependably monitor the B_7_ binding to SAV and presumably to AV. In addition, the absence of any detectable intermediate in the association reaction in both cases is remarkable, since we monitored the initial binding of B_7_ and SAV using the fluorescence change and fluorescence anisotropy signals, and the independent tryptophan-quenching study the final docking of B_7_ near the Trp. Conversely, there is another independent Surface Plasma Resonance (SPR) study of immobilized B_7_ binding to SAV that yielded a slower k_on_ of 5.13 × 10^6^ M^−1^s^−1^ at 4°C,^63^ which was ∼5X slower than our 2.6 × 10^7^ M^−1^s^−1^ at 4 °C, calculated by an Arrhenius plot (ln *k*_on_ vs 1/T) of the BFl data. Similarly to AV, we believe that SPR methodology for the B_7_ and AV-like protein kinetics^20,36^ are modified by the immobilization of one reactant, either B_7_ or protein, to the chip.

### 3.4 Biotinylated-duplex association reaction to AV and SAV

#### 3.4.1 The association rate constants of biotin attached to our biotin-14mer_ds_*Fl

In case of biotinylated 14mer duplex showed a biphasic behavior with two temperature and concentration dependent rate constants (Table 2, Figure 7) when reacting with both AV and SAV. The biphasic association rate constants, k_on1_ and k_on2_, summed to approximately 70 % of the total reaction amplitude. The remaining ∼30% was assigned to a third rate constant (0.02 ± 0.01 s^−1^) that presented neither temperature nor concentration dependence; therefore, it has been assigned to the readjustments of the Fl dye after being displaced by both proteins. The k_on1_ and k_on2_ association rate constants of SAV were 3.4X and 1.8X faster than the corresponding rate constants of AV (Figure 8) as observed with the BFl and BcO probes, confirming the differences in the AV and SAV binding pockets.

**Figure 8.**
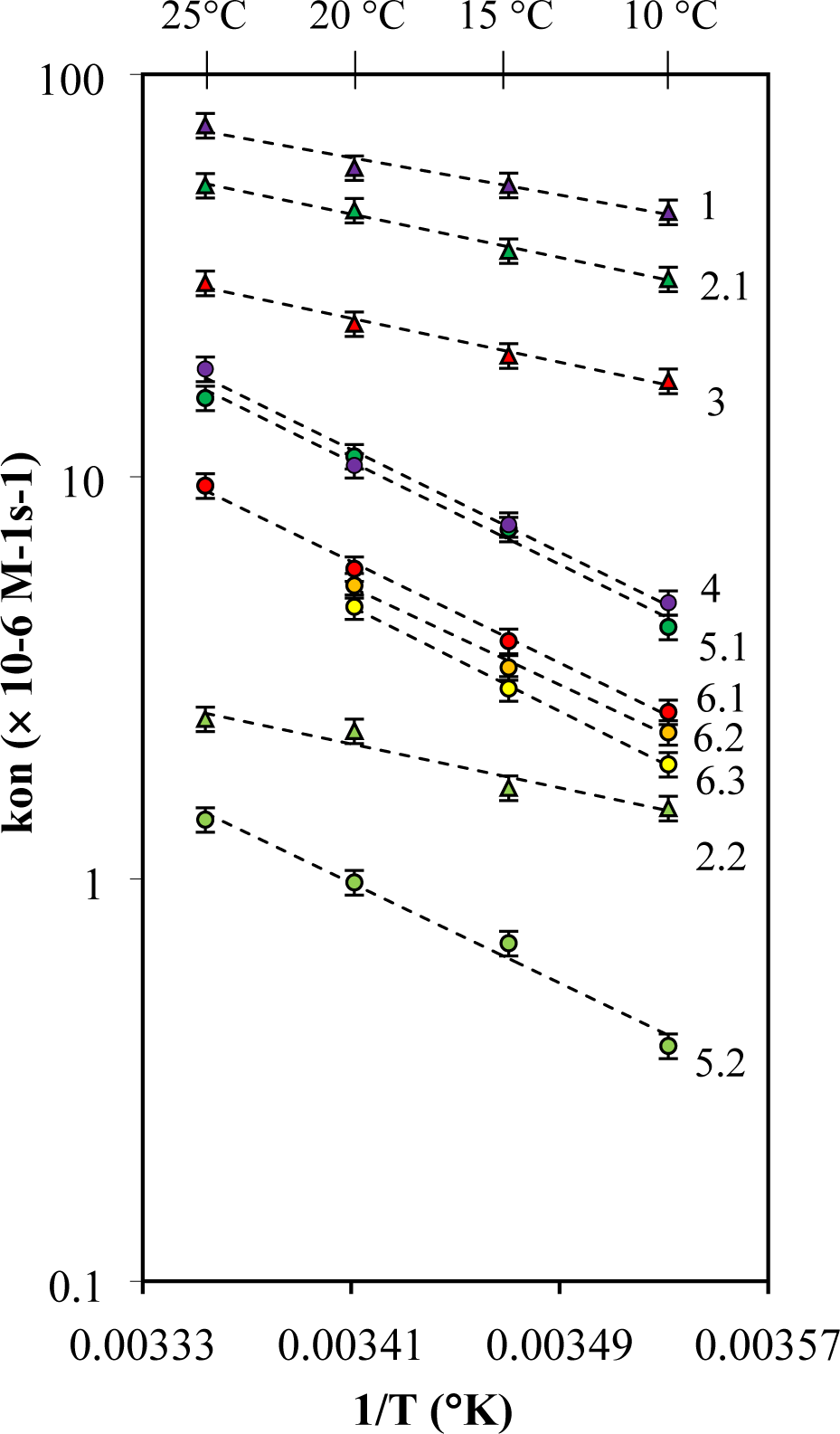
Arrhenius plot (ln k_on_ vs 1/T) of the association rate constants plotted in semi-logarithm for clarity. Temperature dependence of the biotin association reaction at pH 8 (unless otherwise specified) for: 1 SAV-BFl (purple triangles); 2 SAV-biotin-DNA_ds_*Fl (green triangles): 2.1 (k_on1_) and 2.2 (k_on2_); 3 SAV-BcO (red triangles); 4 AV-BFl (purple circles); 5 AV-biotin-DNA_ds_*Fl (green circles): 5.1 (k_on1_) and 5.2 (k_on2_); 6 AV-BcO (red circles): 6.1 at pH 8, 6.2 at pH 9 (orange circles), 6.3 at pH 10 (yellow circles).

**Figure 7.**
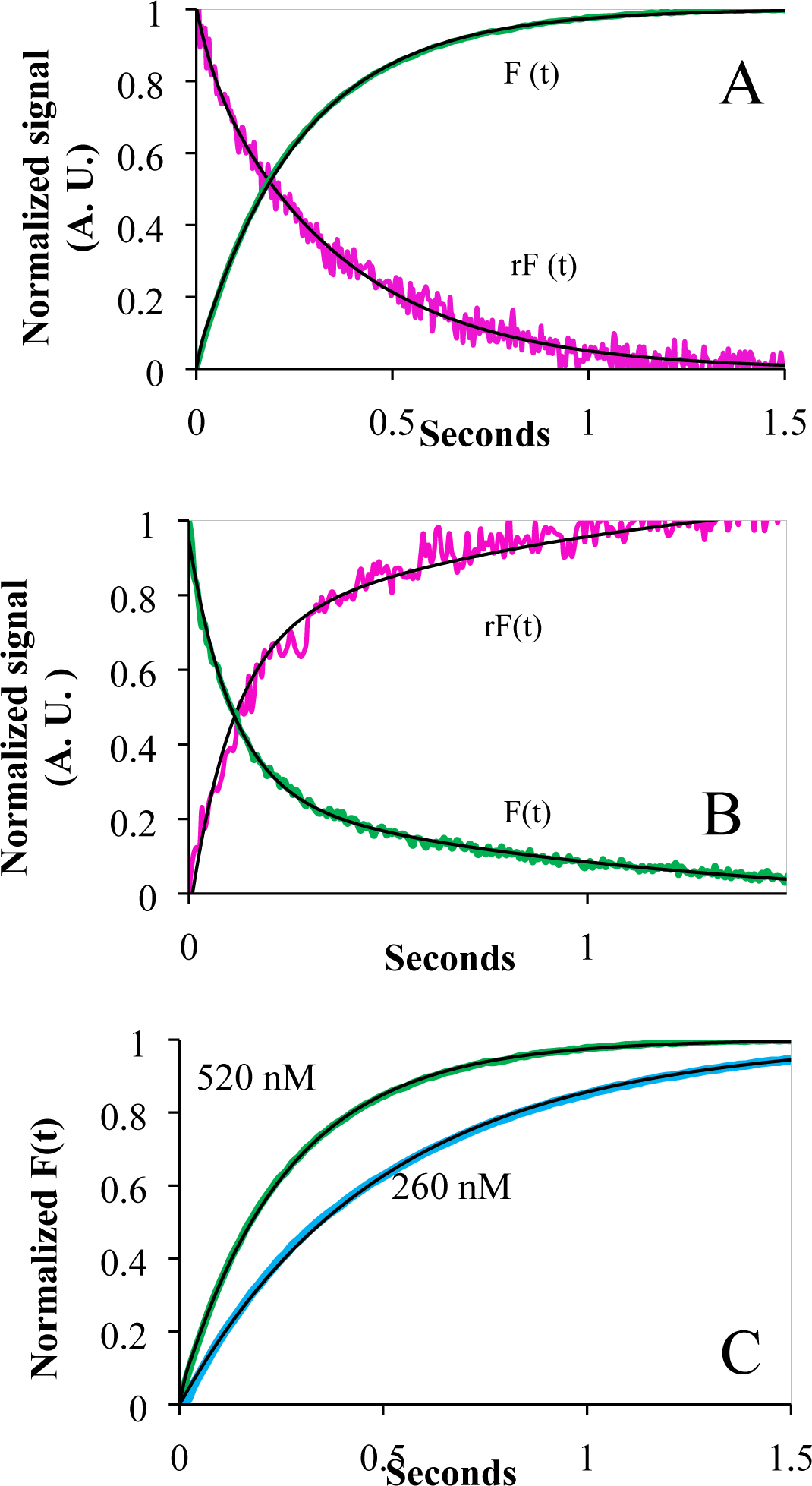
**A)** Biotin-DNA_ds_*Fl (20 nM) binding to: **A)** AV (520 nM) and **B)** SAV (200 nM), at 15 °C. **C)** Concentration dependence of biotin-DNA_ds_*Fl (20 nM) binding to AV at 15 °C. All curves (black line) were strongly biphasic. Notice the inversion of SF signals. However, the *F*(*t*) traces were in prefect agreement with QY experiments.

#### 3.4.2 Comparisons with other biotinylated DNA kinetic studies

An independent FRET study monitored the reaction of B_7_ attached to the 5’ end of a 46 nucleotide duplex DNA binding to SAV.^35^ The reaction also showed two rate constants at pH 8, but at unspecified temperature, pre-exponentials and errors. To make a comparison, we have chosen SAV data at 20 °C whose association rate constant, k_on1_, of 4.59 ± 0.8 × 10^7^ M^−1^s^−1^ was in excellent agreement with the 4.5 × 10^7^ M^−1^s^−1^ reported by the mentioned study. In the case of our k_on2_ of 2.33 ± 0.1 × 10^6^ M^−1^s^−1^, it was in good agreement with the second rate of 3.0 × 10^6^ M^−1^s^−1^ of that independent study. The agreement in the data validates our findings which imply that B_7_ attached internally to DNA (or at the 5’ end) will have two rate constants, one enhanced and other diminished probably due to unfavorable orientation according to the reaction models discussed below.

### 3.5 Significance of the association rate constants

The B_7_ binding to AV and SAV (at 25 °C) were, respectively, between 54-714X and 13-400X slower than 10^9^ M^−1^s^−1^ as expected for a diffusion limited process. ^64^ On the other hand, the k_on_ values of SAV were 3-4X faster than AV’s despite the similarity of the AV and SAV binding sites in the crystal structures (Figure 8). Our deglycosylation experiments indicate that the disparity in the k_on_ values between both SAV and AV proteins cannot be explained by the presence or absence of the carbohydrate motif on the AV but for true collective interactions of the aminoacids in the binding pocket and the biotin ring.

### 3.6 Biotin vs. Biocytin

In our study, the association rates were acquired with B_7_ and Bc probes, BFl and BcO; respectively, in which Biocytin present a longer carbon linker; however, the values only differed by 2-fold (Table 1), from 10 °C to 25 °C, when reacting with AV. It is important to clarify that the association rates were not enhanced by the electrostatic attraction of the negative charged probes (BFl and BcO) and the positive AV;^29^ since, the association rates of those two probes binding to neutral SAV differed also by ∼2 fold as observed for AV. The dissociation constant, K_D_, of AV-B and AV-Bc were reported to be 10^−13^ and 10^−15^ M, respectively, differing by 100-fold.^37^ Consequently, this 100-fold difference, if accurate, must be caused by a difference of 50-fold in the k_off_, dissociation rate constants which is discussed below.

### 3.7 Dissociation kinetics

The dissociation reactions of the AV-B and SAV-B complexes had been described as passive unimolecular “replacements” 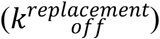 with units of reciprocal seconds (s^−1^) and values of 9 × 10^−8^ s^−1 32^ and 2.4 × 10^−6^ s^−1^,^65^ respectively. However, we have also observed a bimolecular “displacement” off-rate constants 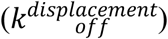 with M-1s-1 units for the SAV-BcO complexes (AB_1_ and AB_4_) that were strongly dependent of B_7_ concentration (Figure 9A) and temperature (Figure 9B). These reactions had ∼79% of the total release amplitude, in contrast to the 5% when BFl was used (Figure 9C); therefore, the longer “tail” of the BcO facilitated the displacement, a feature that can be exploited to increase purification yields. Also, the process is protein dependent, as it was not observed for the AV complexes. Significantly, this new information can find important applications in affinity chromatography purification based on SAV and longer “tail” or tethers that will help to increase the release of the product and enhance efficiency.

### 3.8 Biotin reaction models to AV and SAV

#### 3.8.1 Reaction model of BFl and BcO binding to AV and SAV

The SF traces of B_7_ binding to AV and SAV were best fitted by a simple association model, A + B ⇌C. A single rate constant, k_on_ (Eqn. 8), could be fit with no intermediates or evidence of cooperativity considering that the dissociation reaction was not significant for the first 5-8 sec after mixing. More elaborate mechanism have been reported^66,67^. For example, A + B ⇌C⇌D has been proposed for polystyrene SAV coated particles (6.5 nM) reacting with a fluorescein labeled biotin probe (1.75 nM and 17.5 nM), whose linker resembles our BcO probe. This model required fitting of two dissociation and two association rate constants with the extra equilibrium attributed to two reasons: 1) The interference of the dye structures into the neighboring site due to multiple occupancies on the tetramer^58^ and 2) to possible inhibitory steric interactions caused by high density of SAV sites on the surface of the polystyrene particles. Interestingly, a similar model was used to analyze a pull-off study carried out by Scanning Force Microscopy for AV-B complex with immobilized AV in which two events of 20-40 pico-newtons and 40-80 pico-newtons were assigned to the presence of an intermediated.^68^ Categorically, we have avoided these experimental complications by following the reaction at pseudo first order to ensure that our probes occupied only one binding site of AV and SAV in solution (non-immobilized), as discussed above. However, when considering a particular AV or SAV bioassay, one must consider that surface matrix complexity and multiple orientation of the B_7_ and AV-like proteins can modify the dissociation mechanism with respect to those observed in solution by us.

**Figure 9.**
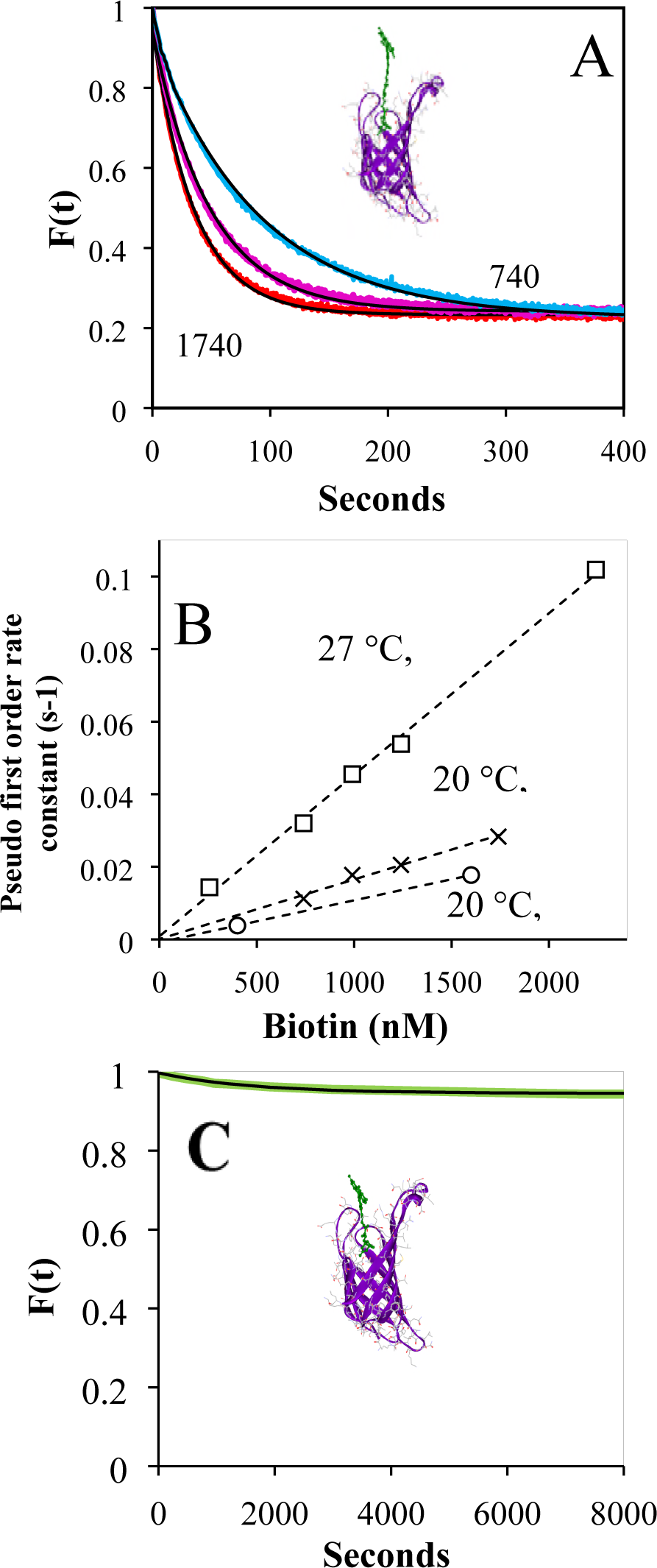
**A)** Concentration dependence of the displacement reaction of SAV-BcO complex (AB_1_ model) by unlabeled biotin at 20 °C. The concentration of challenging B_7_ was 740 nM (blue), 1240 nM (pink) and 1740 nM (red) after the remaining free binding sites were filled. The half-times were 56.6 s, 33.9 s and 24.2 s, respectively; with a release amplitude of 79 ± 1 %. **B)** Temperature dependence of the displacement reaction of SAV-BcO by unlabeled B_7_ for the AB_1_ filling model (at 20 °C and 27 °C) and for the AB_4_ model (at 20 °C). The corresponding 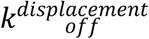 (calculated from the slope) were 1.64 ± 0.4 × 10^5^ M^−1^s^−1^, 4.62 ± 0.3 × 10^5^ M^−1^s^−1^ and 1.20 ± 0.3 × 10^5^ M^−1^s^−1^, respective. **C)** Displacement reaction of unlabeled biotin and SAV-BFl complex (AB_1_ filling model) at 30°C. The concentration of challenging biotin was 1400 nM which produced a release of only 5 % of the bound probe. The green curves is the observed data and black curve is the fitted curve for which only 6.5% displacement was observed for SAV-BFl complex in contrast with 79% in case of the complex formed with the longer linker BcO.

#### 3.8.2. Reaction model of biotin-DNA_ds_*Fl binding to AV and SAV

was best described by two parallel reactions (Eqn. 9) with two independent association rate constants that showed no evidence of intermediates in solution and whose pre-exponentials were temperature dependent (Table 2) indicating the presence of two B_7_ populations with different orientations with respect to the DNA and responsible for the measured *k*_*on1*_ and *k*_*on2*_ rate constants. Thus, at 25 °C, the measured value of *k*_*on1*_ for both AV and SAV were 20-40 % slower than rate constants acquired with BFl indicating that biotin on the DNA was positioned in a favorable orientation that enhance the reaction. On the other hand, the slower k_on2_ rate constant is associated with an unfavorable orientation of the second B_7_ population.

### 3.9. Thermodynamic Parameters

The forward activation energies (E_a_^forward^ or ΔH^‡,forward^) of the B_7_ binding to AV and SAV were ∼6.0 and ∼14 kcal/mol, respectively; and they were in good agreement with early estimation of 10 - 12 kcal/mol for the displacement of water molecules from the binding pocket.^57^ These values were larger than the 3-4 kcal/mol^32,69^ characteristic of a diffusion limited reaction (which requires also association rate constants in the order of 10^9^ M^−1^s^−1^ and our fastest values were in the order of ∼1.9×10^7^ M^−1^s^−1^ and ∼7.5×10^7^ M^−1^s^−1^, at 25 °C for AV-BFl and SAV-BFl, respectively). Hence, the association reaction is not diffusion controlled in the range of experimental work carried by us. Interestingly, the B_7_ binding process for both proteins share the same k_on_ at 52.1 °C (calculated by Arrhenius plot), and mainly that the binding of B_7_ ligand enhances thermal stability of the proteins shifting from 75 °C to 112 °C for SAV and from 84 °C to 117 °C for AV.^70^

Remarkably, the difference of forward and reverse activation energies 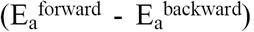, calculated with Arrhenius plots of the association and dissociation rate constants, respectively; matched, within the error, the reaction enthalpy (ΔH°_Rxn_) calculated by calorimetry (Table 3, Figure 10 A). The same argument holds for entropy (ΔS^‡, forward^-ΔS^‡, backwards^) and Gibbs free energy (ΔG^‡, forward^-ΔG^‡, backwards^) and the calorimetric ΔS°_Rxn_ and ΔG°_Rxn_ values calculated by others (see references in Table 3, Figure 10 B, C); Thus, the forward thermodynamic parameters obtained in this study completed nicely the thermodynamics cycles, thus making very compelling arguments in favor of the proposed simple reaction model (Eqn. 8), which has single transition state (^‡^) but no intermediate. The positive nature of ΔE_a_^forward^ and ΔS^‡, forward^ toward the transition state can be explained as the energy required to remove water molecules and displace the protein’s β3-β4 loop^29,71^ with an increment of the overall disorder, ΔS^‡^. A comparative analysis of the transition state (^‡^) for the AV-B and SAV-B complexes reveals that the former has a larger ΔE_a_^forward^ and ΔS^‡,forward^ (Table 3, Figure 10, red line) than the latter (Table 3, Figure 10, green line) which implies that binding sites of AV are deeper and less accessible resulting in a slower association rate constants, k_on_, and larger activation energy with respect to biotin binding to SAV.

**Table 3.**
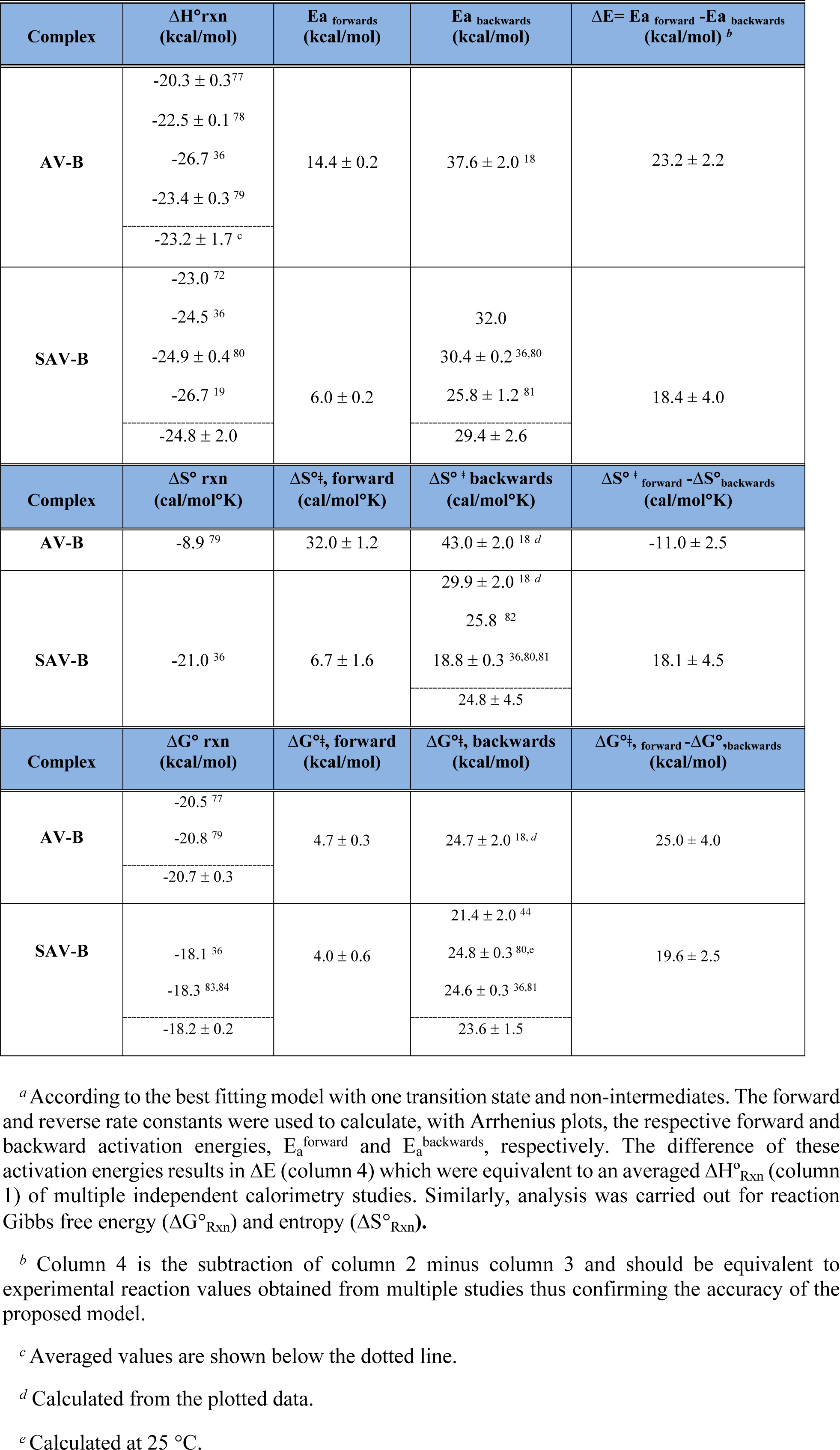
Thermodynamic cycles of biotin binding to AV and SAV for one transition state.^a^

**Figure 10.**
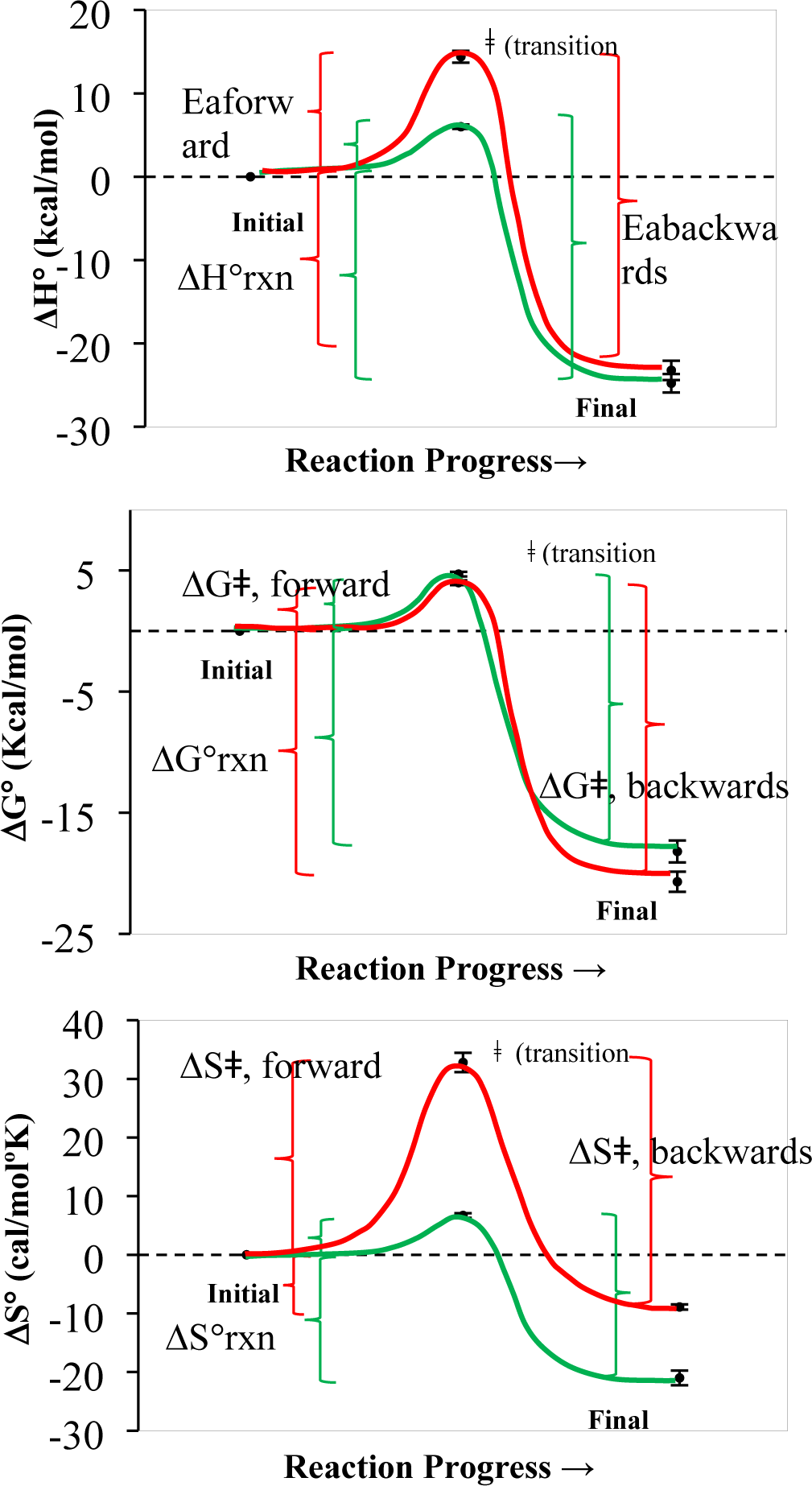
Thermodynamic cycle of the B_7_ binding to AV (red) and SAV (green) for one transition state and no intermediate: **A)** ΔH°_rxn_, **B)** ΔS°_rxn_ and **C)** ΔG°_rxn_ is the average of values found in multiple studies (Table 3). Arrhenius plots of the temperature dependent association and dissociation rate constants were used to calculate the E_a_^forward^ and E_a_^backwards^, respectively.

#### The fluorescence spectroscopic parameters

The absorbance and emission peaks of all the dye-labeled B_7_ complexes (Table 4) were red shifted a few nanometers (Supporting Information, Figure S1 and S2) with respect to the unbound probes, with the exception of the biotin-DNA_ds_*Fl complexes with AV and SAV that were blue-shifted 3 nm by the presence of both proteins. This can be explained due to fluorescein (Fl) interactions with DNA_ds_ before binding to AV and SAV and later displaced toward the solution in the complex. In the particular case of the absorbance spectrum of SAV-BFl, it was highly distorted (Supporting Information, Figure S1 B) owing to the shifting of the Fl^−2^/Fl^−1^ equilibrium by charge transfer;^72^ since, we detected the corresponding 4.1 ns and 3.0 ns lifetimes (τ). The time-resolved fluorescence of biotin-DNA_ds_*Fl complexes of both proteins (Supporting Information, Table S3) had two lifetimes decays whose exponentials were not affected by temperature suggesting the existence of the two Fl positions on the DNA which make a compelling argument for the parallel reaction model (Eqn. 9) with two reacting populations: (*Biotin-DNA*_*ds*_ ^*\**^ *Fl*)_1_ and (*Biotin-DNA*_*ds*_ ^*\**^ *Fl*)_2_,

**Table 4.**
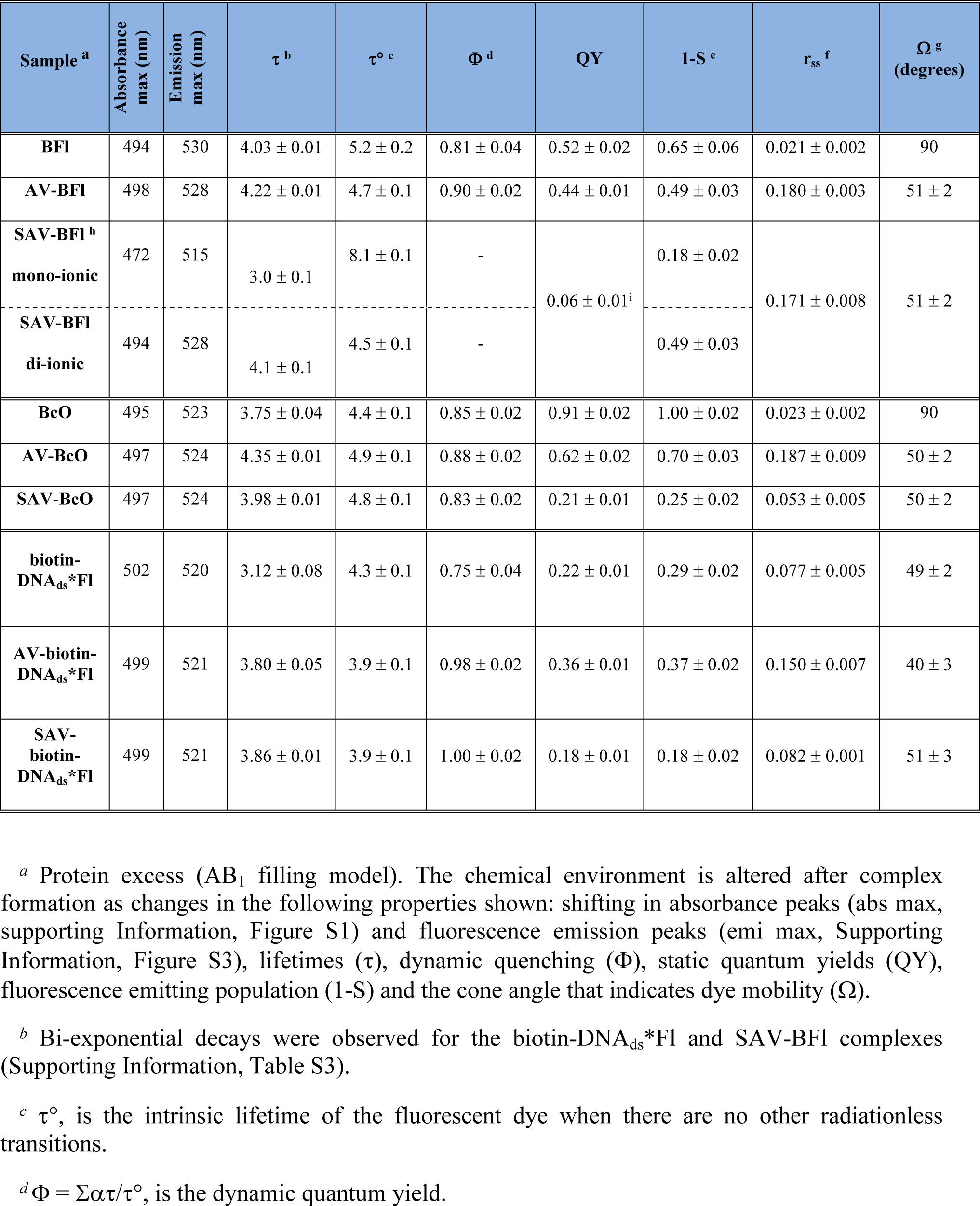
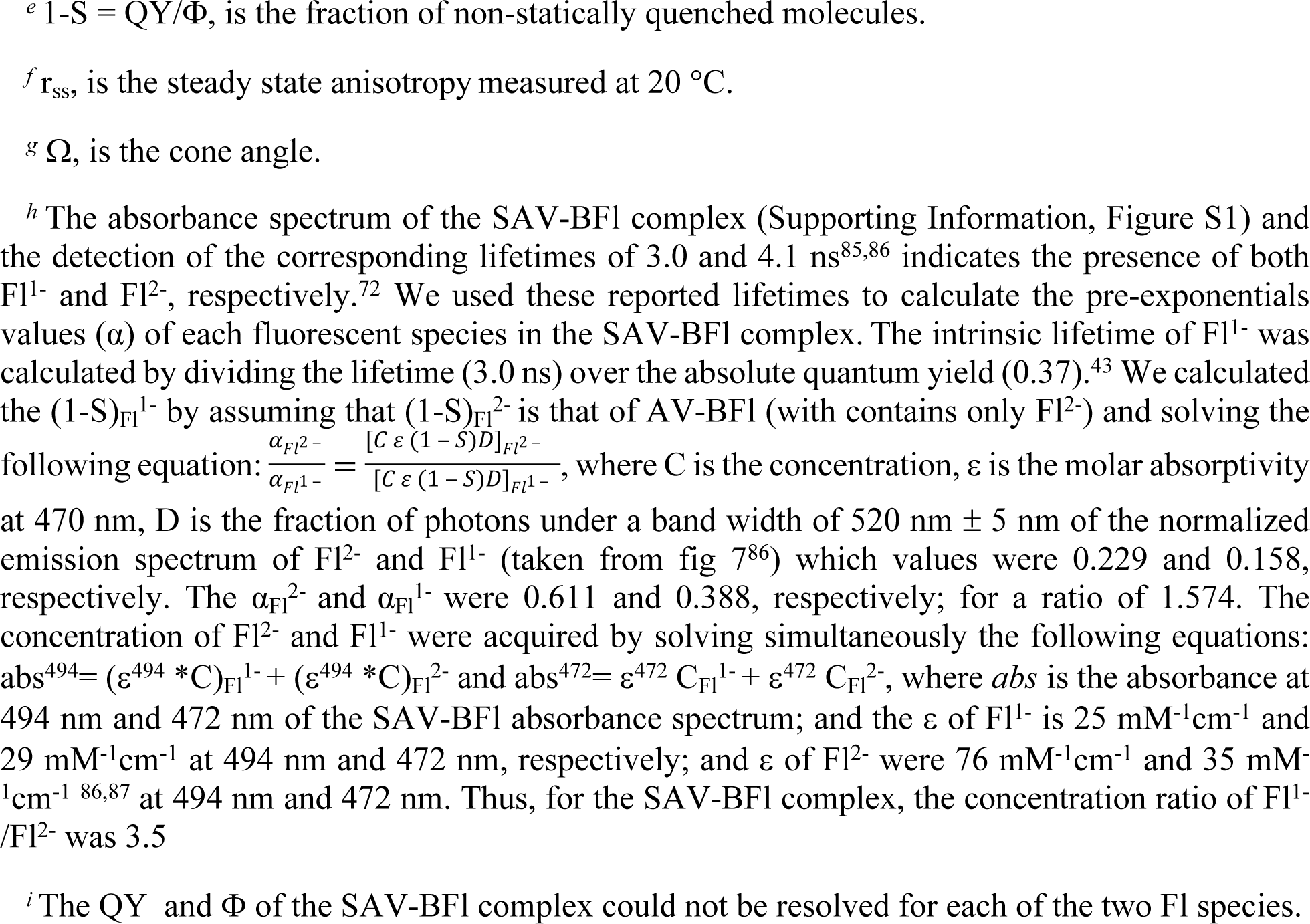
Spectroscopic parameters of the dye-labeled biotin probes and respective complexes with AV and SAV.

| Sample <sup>a</sup> | Absorbance max (nm) | Emission max (nm) | $\tau$ <sup>b</sup> | $\tau^{\circ}$ <sup>c</sup> | $\Phi$ <sup>d</sup> | QY | 1-S <sup>e</sup> | $r_{ss}$ <sup>f</sup> | $\Omega$ <sup>g</sup> (degrees) |
| --- | --- | --- | --- | --- | --- | --- | --- | --- | --- |
| **BFI** | 494 | 530 | $4.03 \pm 0.01$ | $5.2 \pm 0.2$ | $0.81 \pm 0.04$ | $0.52 \pm 0.02$ | $0.65 \pm 0.06$ | $0.021 \pm 0.002$ | 90 |
| **AV-BFI** | 498 | 528 | $4.22 \pm 0.01$ | $4.7 \pm 0.1$ | $0.90 \pm 0.02$ | $0.44 \pm 0.01$ | $0.49 \pm 0.03$ | $0.180 \pm 0.003$ | $51 \pm 2$ |
| **SAV-BFI <sup>h</sup>**<br>**mono-ionic** | 472 | 515 | $3.0 \pm 0.1$ | $8.1 \pm 0.1$ | - | $0.06 \pm 0.01^i$ | $0.18 \pm 0.02$ | $0.171 \pm 0.008$ | $51 \pm 2$ |
| **SAV-BFI**<br>**di-ionic** | 494 | 528 | $4.1 \pm 0.1$ | $4.5 \pm 0.1$ | - | | $0.49 \pm 0.03$ | | |
| **BcO** | 495 | 523 | $3.75 \pm 0.04$ | $4.4 \pm 0.1$ | $0.85 \pm 0.02$ | $0.91 \pm 0.02$ | $1.00 \pm 0.02$ | $0.023 \pm 0.002$ | 90 |
| **AV-BcO** | 497 | 524 | $4.35 \pm 0.01$ | $4.9 \pm 0.1$ | $0.88 \pm 0.02$ | $0.62 \pm 0.02$ | $0.70 \pm 0.03$ | $0.187 \pm 0.009$ | $50 \pm 2$ |
| **SAV-BcO** | 497 | 524 | $3.98 \pm 0.01$ | $4.8 \pm 0.1$ | $0.83 \pm 0.02$ | $0.21 \pm 0.01$ | $0.25 \pm 0.02$ | $0.053 \pm 0.005$ | $50 \pm 2$ |
| **biotin-DNA<sub>ds</sub>\*FI** | 502 | 520 | $3.12 \pm 0.08$ | $4.3 \pm 0.1$ | $0.75 \pm 0.04$ | $0.22 \pm 0.01$ | $0.29 \pm 0.02$ | $0.077 \pm 0.005$ | $49 \pm 2$ |
| **AV-biotin-DNA<sub>ds</sub>\*FI** | 499 | 521 | $3.80 \pm 0.05$ | $3.9 \pm 0.1$ | $0.98 \pm 0.02$ | $0.36 \pm 0.01$ | $0.37 \pm 0.02$ | $0.150 \pm 0.007$ | $40 \pm 3$ |
| **SAV-biotin-DNA<sub>ds</sub>\*FI** | 499 | 521 | $3.86 \pm 0.01$ | $3.9 \pm 0.1$ | $1.00 \pm 0.02$ | $0.18 \pm 0.01$ | $0.18 \pm 0.02$ | $0.082 \pm 0.001$ | $51 \pm 3$ |

<sup>a</sup> Protein excess (AB<sub>1</sub> filling model). The chemical environment is altered after complex formation as changes in the following properties shown: shifting in absorbance peaks (abs max, supporting Information, Figure S1) and fluorescence emission peaks (emi max, Supporting Information, Figure S3), lifetimes ( $\tau$ ), dynamic quenching ( $\Phi$ ), static quantum yields (QY), fluorescence emitting population (1-S) and the cone angle that indicates dye mobility ( $\Omega$ ).
<sup>b</sup> Bi-exponential decays were observed for the biotin-DNA<sub>ds</sub>\*FI and SAV-BFI complexes (Supporting Information, Table S3).
<sup>c</sup> $\tau^{\circ}$ , is the intrinsic lifetime of the fluorescent dye when there are no other radiationless transitions.
<sup>d</sup> $\Phi = \Sigma \alpha \tau / \tau^{\circ}$ , is the dynamic quantum yield.
<sup>e</sup> $1-S = QY/\Phi$ , is the fraction of non-statically quenched molecules.

The deconvolution of the SF binding traces was completed using the steady-state anisotropy (r_ss_) whose AV values were larger than SAV attributable to a larger molecular weight and to the presence of the carbohydrate motif for the former. Significantly, the quantum yields (QY) of the complexes were in excellent agreement with all the binding association traces, which is particularly important for the biotin-DNA_ds_*Fl reactions, whose traces had shifted directions (Figure 7) owing to opposite quenching interactions from the initial quantum yield, QY=0.22 ± 0.01, of the free probe biotin-DNA_ds_*Fl (Table 4) to an increment up to 0.36 ± 0.01 for the AV-biotin-DNA_ds_*Fl complex and a decrement to 0.18 ± 0.01 for the SAV-biotin-DNA_ds_*Fl complex. This effect is caused by the bulker nature of the AV with respect to SAV that allows further displacement of Fl from the 3’ end toward the solution environment resulting in the increase of the QY for the biotin-DNA_ds_*Fl-AV.

The (S) and (1-S) are, respectively, the emitting and statically quenched dye populations. The latter always increased with the complex formation with respect to the unbound free probes; thus, the fluorescence information pertained to the self-revealing population whose cone angles (Ω) of ∼50° pointed out that the dye probe was fairly free to rotate (Figure 11) in the complexes. On the other hand, the presence of (1-S) did not affect the accuracy of association rate values, as the rates obtained in the independent SAV tryptophan-quenching study^36^ and our data were in perfect agreement.

**Figure 11.**
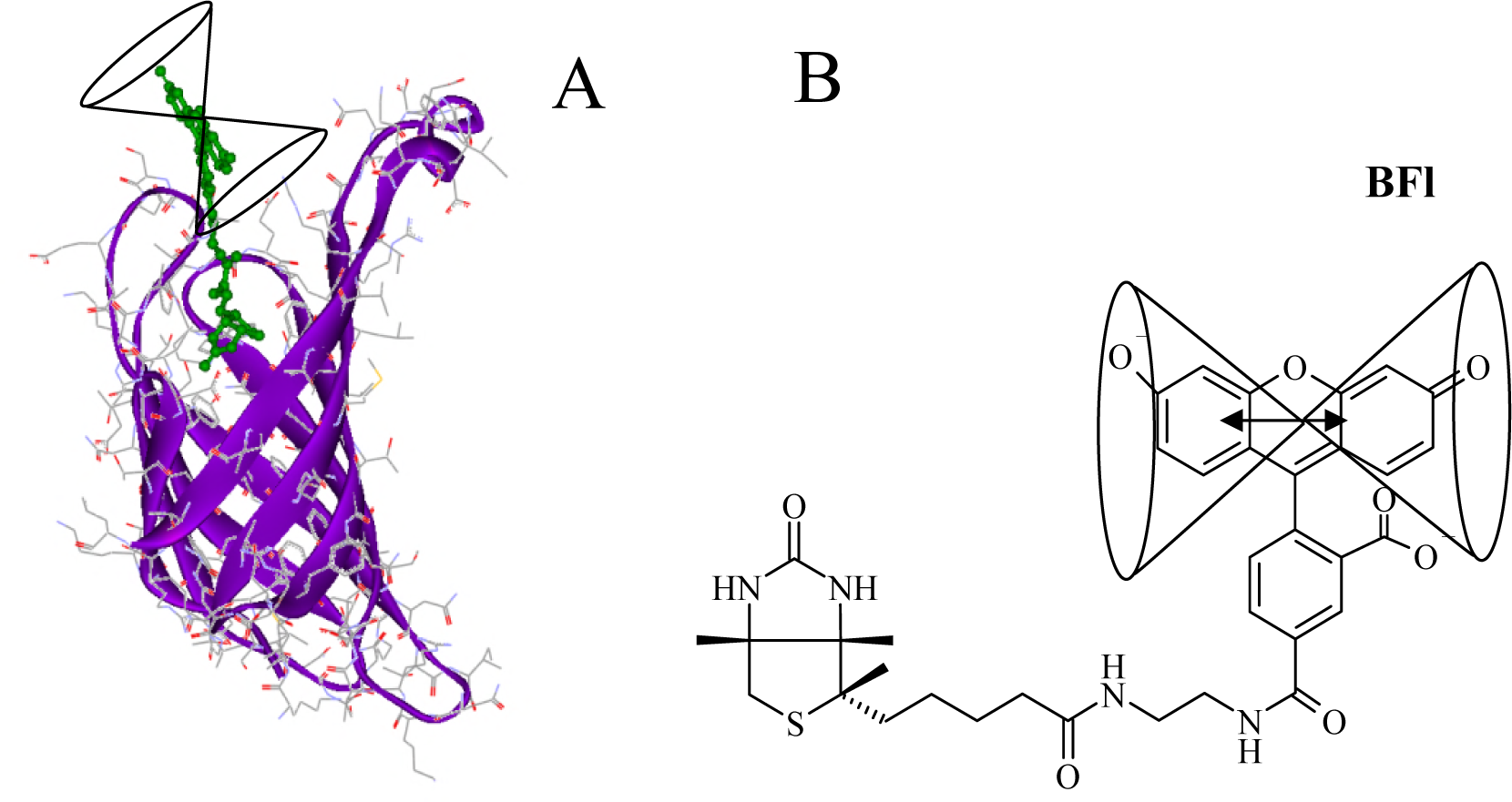
A) Pictorial representation of the AV-BFl complex for only one binding sites showing a relative high dye mobility with a half apical angle^88^ (Ω) of 51° ± 2° in contrast with the unrestricted mobility of the dye labeled B_7_ with 90°. The figures reflect rotational motion of the transition moment of the isoalloxanzine ring system within the cone B). The dye structure added using Accelrys DS visualizer 2.0 ® to the AV-B crystal structure complex: 2AVI.^29^

## 4. CONCLUSIONS

In the presented study, we calculated the association rate constants of B_7_ binding to AV and SAV with dye-labeled B_7_ probes and unlabeled B_7_. We concluded that attached fluorescent probes did not alter the association rates and no binding cooperativity was observed when comparing the initial (unoccupied) and final (occupied) binding rates. The fluorescence, 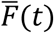, and corrected anisotropy signals, 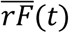, of the dye-labeled B_7_ probes provided truthful binding traces contrary to the uncorrected anisotropy signal, 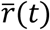, due to changes in the QY of the participating reacting species. The B_7_ association rate constants of SAV are several times faster than AV and the glycan chain of the latter does not play a role in the B_7_ binding association and neither explains the difference in the k_on_ values between these two proteins. Thus, we conclude that the main differences in reaction speeds is likely related to structural aspects of the binding sites. A deeper pocket in AV would be filled with more water molecules requiring more energy to break the hydrogen bonds. Also, the variation in requirements for induced fit could explain in larger activation energy and entropic increment for AV compared to the SAV in the overall thermodynamics of the reaction. Interestingly, the overall reaction free energy changes are equivalent.

The association rate constant for BcO, in which the tag is attached to a longer linker of biocytin, is ∼2X faster than B_7_ with the shorter linker (BFl) for both proteins. The difference of 100X in K_D_ of AV complex with biotin and biocytin can be explained by differences in the dissociation process rather than the association rate constants. The B_7_ binding to AV and SAV is not diffusion limited as larger than 3 kcal/mol activation energies were calculated with Arrhenius plots of the rate constants, and those all of those rates were two orders of magnitude slower (on average ∼10^7^ M^−1^s^−1^) than the 10^9^ M^−1^s^−1^ required for diffusion limited reactions. The forward thermodynamic parameters of B_7_ binding to AV and SAV complemented nicely the thermodynamic cycles with data obtained with independent calorimetric studies and dissociation kinetics elsewhere. Thus, the most probable reaction model is the one with non-intermediate and a single transition state in solution but it could be more elaborate on support matrices, such as chips.

The spectroscopic properties indicated very compact complexes with high dye mobility for all the probes, BFl, BcO and biotin-DNA_ds_*Fl. We report for the first time a bimolecular displacement process for the SAV-BcO complex when challenged by unlabeled B_7_ and this displacement rate constant for the B_7_ with the longer linker (biocytin) in the BcO suggests that repair and recondition of enriched biotin-avidin-like surfaces is possible if even long linkers are used. A potential application of dye-labeled B_7_ and AV-like complex could be in Dye-Sensitized Solar Cells (DSSC)^73,74^ as the AV and SAV complexes are highly thermally stable at 112 °C and 117 °C^70^, respectively, may perhaps be attached covalently to the n-type material (e.g. TiO_2_) and the charge-transfer molecule to B; e.g. porphyrins, chlorophylls, ruthenium-complexes, coumarins or indoline dyes, ^75^ with the advantage of regeneration capabilities, as damage dye can be reconditioned or replaced by another dye type on the tetramer attached surface (Figure 9A). The spectroscopic properties of these dye-labeled B_7_ and AV-like complexes are vital for detection methods based on polarization, fluorescence, anisotropy and Fluorescence Resonance Energy Transfer (FRET) systems because static, dynamic quenching and rotational constrains of the fluorescent probes reduce the detection limits by decreasing the signal to noise ratios. ^76^ and producing artifacts. The information in here presented will be valuable to improve new nano-technological applications of B_7_ and AV-like protein systems.

## ASSOCIATED CONTENT

**Supporting Information Available**: “Displacement reaction of SAV-BFl by unlabeled B, absorbance and emission spectra of dye-labeled B_7_ complexes, and fluorescence lifetime table”. The following files are available free of charge.

**Figure S1.** Absorbance spectra of dye-labeled biotin probes (PDF).

**Figure S2.** Fluorescence emission spectra of dye-labeled biotin probes (PDF).

**Table S3.** Lifetimes of dye-labeled biotin probes and protein complexes (PDF).

## Funding Sources

The work was supported by: National Institutes of Health Grants GM59346 and RR015468 to LJP; CONACYT-Mexico postdoctoral, SNI fellowships (130994, 162809, SNI75487) and the Government of Veracruz-Mexico gifted-student fellowships to RFD; Bioengineering, Biosystems and Synthetic Biology Focus Group of Tecnológico de Monterrey (002EICIP01).

### ACKNOWLEDGMENT

This work was carried out at Department of Chemistry, University of Nebraska-Lincoln, NE 68588-0304, USA. Roberto F Delgadillo thanks Dr. Efrain Barragan, Dr. Omar Olmos-Lopez and Ms. Lola R. for their support; Andrea R. Gomez-Fernandez and Yuriana Oropeza for proof reading the manuscript; CONACYT-Mexico for the postdoctoral and SNI grants; and the Government of Veracruz-Mexico. Roberto F. Delgadillo and José González-Valdez would like to thank the Bioengineering, Biosystems and Synthetic Biology Focus Group of Tecnológico de Monterrey (002EICIP01) for its support.

**Author Contributions:** The manuscript was written through contributions of all authors. All authors have given approval to the final version of the manuscript.

